# High-yield protein production in the chemolithoautotrophic bacterium *Cupriavidus necator* H16

**DOI:** 10.1101/2025.11.17.688765

**Authors:** Matteo Vajente, Marleen Hallamaa, Hendrik Ballerstedt, Lars Mathias Blank, Sandy Schmidt

## Abstract

Many chemical manufacturing routes are being replaced with enzymatic processes to improve sustainability. In biocatalysis, enzyme production is often the main bottleneck. Bacterial proteins are frequently produced using the workhorse *E. coli* BL21(DE3) and its derivatives. However, other bacteria with beneficial characteristics can also be engineered for this purpose. *Cupriavidus necator* H16 (*C. necator*), for example, is a Gram-negative bacterium well-known for its chemolithoautotrophic metabolism and high polyhydroxybutyrate (PHB) accumulation. Previous studies have demonstrated protein production without inclusion body formation, which is one of the main challenges when producing enzymes in *E. coli*. Nevertheless, high-yield protein production in *C. necator* remains an understudied field. Here, we investigated the bottlenecks limiting protein production in *C. necator*. We optimized a T7 RNA polymerase genetic system and quantified the impact of several genetic elements such as T7 promoter, RBS strength and codon usage towards GFP production. Codon usage was the main factor limiting protein production in *C. necator*. Tuning the RBS strength and selecting a different T7 promoter strongly influenced expression leakiness. We then produced the ene-reductase YqjM from *Bacillus subtilis* in *C. necator*, analyzing the performance of the engineered *C. necator* strain and our previously developed pMVRha expression plasmid. *C. necator* produced high amounts of soluble protein and outperformed the gold standard, *E. coli* BL21(DE3) in producing FMN-loaded enzyme. This result highlights the potential of non-model bacteria to achieve high-yield enzyme production and promote the transition to biocatalysis-driven chemical synthesis.

**Graphical Abstract:** 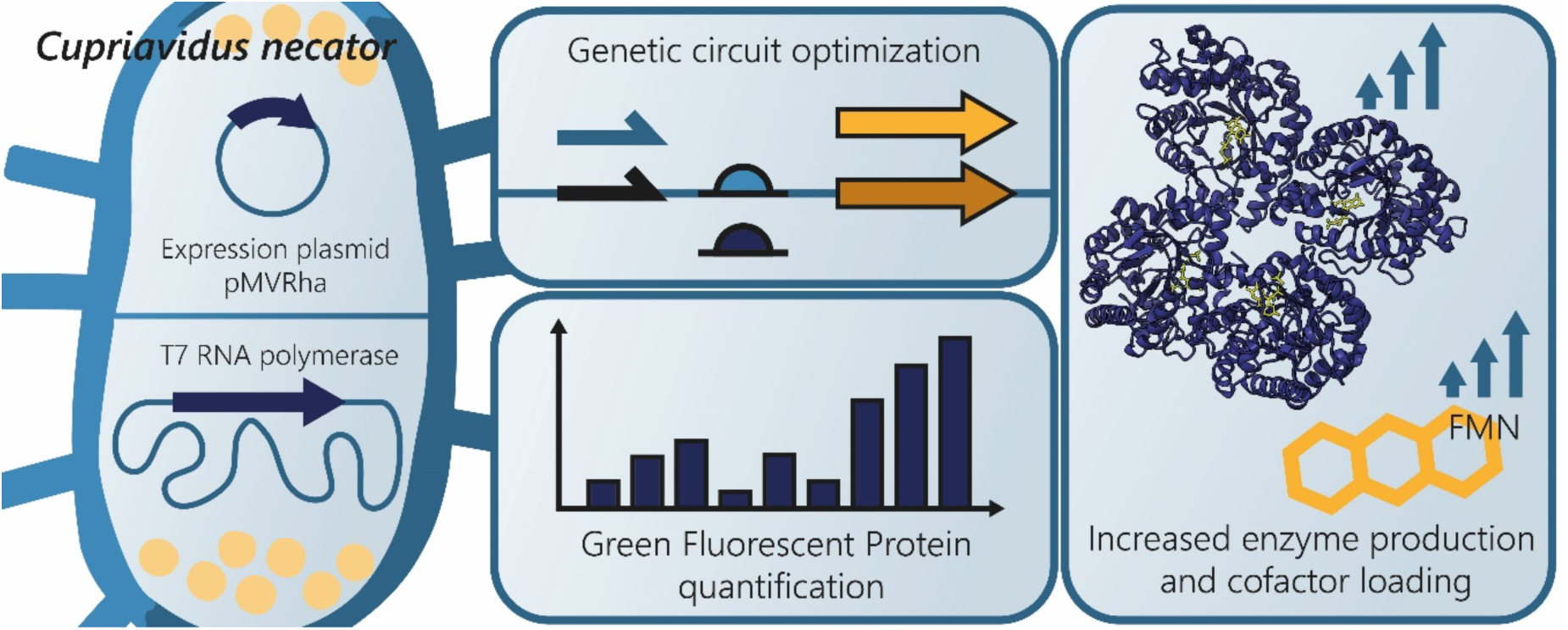

## 1. Introduction

Biocatalysis has emerged as a cornerstone of biotechnology, offering efficient alternatives to traditional chemical synthesis by harnessing the high selectivity and reactivity of enzymes. Central to advancing biocatalysis is the reliable and cost-effective production of enzymes, which requires not only optimization of the specific protein but also careful selection of the host system in which it is produced. Different prokaryotic and eukaryotic hosts offer distinct advantages in terms of expression yield, post-translational modifications, scalability, and economic feasibility (Tripathi and Shrivastava, 2019). Since the first application in 1986, *Escherichia coli* (*E. coli*) strains from the B lineage containing T7 RNA polymerase (T7RNAP) have become the gold standard for protein production in prokaryotic organisms (Studier and Moffatt, 1986). The T7 system allows easy induction and strong orthogonal expression of specific genes of interest. While novel *E. coli* strains have been developed to address shortcomings of the first system (e.g., leaky expression of the gene of interest) (Rosano et al., 2019), growing *E. coli* to high-cell density still suffers from various drawbacks including the formation of organic acids, proteolysis and inclusion-body formation (Choi et al., 2006; Shiloach and Fass, 2005). Moreover, several heterologous proteins have been troublesome to produce in *E. coli* due to a lack of appropriate chaperones or maturation factors (Ryu et al., 2024; Schiffels et al., 2013). Other non-model bacteria might be more suitable for the task. In particular, autotrophic organisms such as *Cupriavidus necator* (*C. necator*, formerly known as *Ralstonia eutropha*), are currently attracting attention because of their potential applications in C1-based biotechnology. However, efficient protein production is still underdeveloped in such non-model strains. *C. necator* is an attractive host for protein production due to its versatile proteome (Jahn et al., 2021) and its chemolithoautotrophic metabolism. It is able to grow autotrophically on a mixture of H_2_, O_2_, and CO_2_, and engineered strains have shown production of various metabolites from CO_2_ (Pan et al., 2021). Enzyme production has been demonstrated in both heterotrophic and autotrophic conditions, proving that protein production in autotrophic conditions is feasible (Arhar et al., 2024). In addition, the native formatotrophic metabolism of *C. necator* enables growth on formic acid, which can be synthesized from CO_2_ through electroreduction (Jhong et al., 2013). Recently, formatotrophic enzyme production in this host has also been reported (Hallamaa et al., 2025). Therefore, *C. necator* is a suitable chassis for upgrading CO_2_-rich effluents into proteins of interest for more sustainable biochemical synthesis. This microorganism can be grown at high cell densities in both autotrophic and heterotrophic conditions (Srinivasan et al., 2002; Tanaka et al., 1995). Its high protein content, which can account for up to 83% of cell dry weight depending on the carbon source and growth conditions (Ismail et al., 2025, 2024), emphasizes its potential for producing recombinant proteins, either as a whole-cell biocatalyst or for further protein purification. Interestingly, *C. necator* has been shown to express enzymes that *E. coli* is not able to produce or that cause the formation of inclusion bodies (Ryu et al., 2024; Srinivasan et al., 2002). Food proteins are another promising market sector for hydrogen-oxidizing bacteria (Angenent et al., 2022; Bernal-Cabas et al., 2025), as *C. necator* has QPS status (Qualified Presumption of Safety) under the European Food Safety Authority (EFSA) (EFSA BIOHAZ Panel et al., 2026).

Thus far, the best-performing recombinant expression method in *C. necator* used a T7RNAP-based system; however, it relied on a proprietary strain derived from H16 (Barnard et al., 2004; Byrom, 1994). A recent attempt to establish a T7-based system in *C. necator* H16 was unsuccessful, as the T7-driven system yielded the same amount of Red Fluorescent Protein (RFP) as a plasmid carrying the P_BAD_ arabinose-inducible promoter (Hu et al., 2020). An ideal production strain should enable high-level, tightly regulated gene expression, resulting in high yields of soluble, cofactor-loaded enzyme. Tight regulation is particularly important under autotrophic conditions, especially when energy or carbon availability is limited by process parameters. Such a two-step process could also decrease plasmid loss during the biomass accumulation phase, a known problem in *C. necator* (Ehsaan et al., 2021). While extensive engineering has optimized protein production in *E. coli* BL21(DE3) and its derivatives, comparable efforts in *C. necator* remain limited. Although its potential for CO₂-based small-molecule production is well established (Pan et al., 2021), only a few studies have quantified heterologous enzyme production in this host (Arhar et al., 2024; Assil-Companioni et al., 2019; Barnard et al., 2004; Gruber et al., 2016; Ryu et al., 2024).

In this study, we describe the establishment and optimization of an inducible T7 RNA polymerase system in *C. necator*. During several rounds of iterative engineering, we examined the impact of RBS strength and codon usage on the expression of green fluorescent protein (GFP). We investigated different inducible promoters (L-rhamnose and salicylate) and variants of the T7 promoter. Finally, we evaluated the performance of *C. necator* by producing YqjM, a flavin-dependent ene-reductase, using our newly developed *C. necator* T7R2 strain and our previously created pMVRha expression plasmid. All the genetic parts developed in this study are compatible with the MoClo cloning system. This standardization effort, together with the efficient electroporation protocols available for these *C. necator* strains, shortens the strain creation time considerably and makes it an attractive microbial production chassis. The plasmid cross-compatibility also allows plasmid exchange between *E. coli* and *C. necator*, providing a simple option for switching hosts when protein expression proves to be difficult in *E. coli*. This is especially promising given the vast genome and availability of maturation cofactors in *C. necator*. Furthermore, its native chemolithoautotrophic metabolism enables enzyme production directly from CO_2_ or formate, making *C. necator* a promising chassis for C1-based biocatalysis.

## 2. Materials and Methods

### 2.1. Chemicals, Bacterial Strains, and Culture Conditions

All chemicals used were purchased from Sigma−Aldrich Ltd., VWR International LLC, or Carl Roth GmbH in the highest available purity. The strains, plasmids, genetic elements, and gene sequences used in this study are listed in the Supplementary Information. *C. necator* and *E. coli* were grown in *lysogeny* broth (LB, 10 g/L tryptone, 10 g/L NaCl, 5 g/L yeast extract) at 30 °C (*C. necator*) and 37 °C (*E. coli*), shaking orbitally at 200 rpm. Agar agar was added to a final concentration of 2% to obtain LB agar. When appropriate, growth media were supplemented with antibiotics at the specified concentrations: kanamycin (*E. coli*: 50 μg/mL, *C. necator*: 200 μg/mL (routine cultivations) or 400 μg/mL (selection after electroporation)), tetracycline (*E. coli*, *C. necator*: 15 μg/mL), and gentamicin (*C. necator*: 20 μg/mL).

### 2.2. Cloning and plasmid assembly

All plasmids were assembled using Golden Gate assembly (Engler et al., 2008). Briefly, 75 ng of PCR-amplified backbone or acceptor plasmid were mixed with linear fragments in a 1:2 molar ratio or with annealed oligonucleotides in a 1:10 molar ratio. Golden Gate reactions were carried out in a total volume of 10 μL by mixing the DNA, ultrapure water, T4 DNA ligase buffer, T4 DNA ligase (200 U), and BsaI-HFv2 (6 U). The mixture was then incubated in a thermocycler using the following program: (37 °C, 5 min → 16 °C, 5 min) × 15-30 cycles → 60 °C, 5 min. Finally, the assembled plasmids were used to transform competent *E. coli* DH5α cells. The DNA fragments used to assemble each plasmid are described in detail In Table S3. Schematic diagrams for the cloning of all plasmids are provided in Figures S2−S4. To anneal oligonucleotides, equimolar amounts of complementary primers (final concentration: 30 μM) were mixed in annealing buffer (10 mM Tris, 50 mM NaCl, 1 mM EDTA, pH 7.5) and incubated in a thermocycler for 2 min at 95 °C. The temperature was then decreased by 1 °C per cycle for 70 cycles (40 s each).

### 2.3. Analysis of the codon usage bias in *C. necator* and *E. coli*

The genomic coding sequences from *E. coli* and *C. necator* were retrieved from NCBI (Accession numbers in Table S4). Codon usage tables were calculated from the genomic coding sequences on the Galaxy web platform, using cusp (EMBOSS) on the public server at usegalaxy.eu (Rice et al., 2000; The Galaxy Community et al., 2024). The codons were then grouped based on their relative abundance (“Fraction” variable returned by cusp) and plotted as a histogram.

### 2.4. Codon harmonization and Codon Adaptation Index analysis

T7RNAP and YqjM were codon harmonized for *C. necator* using a previously reported codon-harmonization tool (Claassens et al., 2017). Codon Adaptation Index (CAI) values were calculated using cai custom (EMBOSS) on usegalaxy.eu (Rice et al., 2000; The Galaxy Community et al., 2024). Nucleic sequences and CAI values for all genes used in this manuscript are available in the Supplementary Information (Tables S3, S5). *GFPmut3* was codon optimized, because codon harmonization was impossible (the genome of the original organism was unavailable at the time) using the codon-optimization tool from Geneious Prime (Dotmatics) and the *C. necator* codon usage table calculated as described above.

### 2.5. *C. necator* electroporation and gene manipulation

Electroporation of *C. necator* was performed as previously described (Vajente et al., 2024). Briefly, a single colony of *C. necator* was cultivated in SOB media (950 μL; 20 g/L tryptone, 5 g/L yeast extract, 0.5 g/L NaCl, 2.5 mM KCl, 20 mM MgSO_4_, pH 7) supplemented with 20 μg/mL gentamicin for 16 h at 30 °C. Fresh SOB supplemented with gentamicin was inoculated with the preculture at an initial OD_600_ of 0.1 and cultivated at 30 °C. When the cells reached an OD_600_ of 0.6, they were transferred onto ice and chilled for 5−10 min. The cells were then transferred to 50 mL tubes and centrifuged at 6000 x *g* at 4 °C for 2 min. The supernatant was removed, and the cells were resuspended in 25 mL of ice-cold 50 mM CaCl_2_ by briefly vortexing. They were then incubated on ice for 15 min. The cells were then centrifuged at 6500 x *g* at 4 °C for 2 min, and the supernatant was removed. Cells were washed twice using 25 and 15 mL of ice-cold 0.2 M sucrose, respectively. At the end of each wash, cells were centrifuged at 6,500 x *g* at 4 °C for 3 min, and the supernatant was decanted. The cell pellet was finally resuspended in 1/100 of the initial volume (e.g., 100 mL initial cell culture to 1 mL final resuspension volume). Aliquots of competent cells were frozen in liquid nitrogen and stored at −80 °C until use. For electroporation, each aliquot was thawed on ice for 20 min, transferred into a pre-chilled 1 mm electroporation cuvette, mixed with 50−200 ng of plasmid DNA and electroporated (25 μF, 200 Ω, 1.15 kV). SOB supplemented with fructose (20 mM) was immediately added, and the cells were transferred to a 2 mL centrifuge tube for outgrowth at 30 °C for 2 h. After the outgrowth, cells were diluted and plated on appropriate selective media.

Knock-in and knock-out plasmids were used for *C. necator* H16 genome modification by double homologous recombination, as explained previously (Vajente et al., 2024). When selecting for *ΔnagR* strains (*C. necator ΔnagR*, T7R, T7S), different dilutions were plated on minimum media supplemented with glucose.

### 2.6. Fluorescence Measurement

Precultures were cultivated overnight at 30 °C at 200 rpm in 96-deep well plate wells (Greiner, Masterblock, 96 well, 2 mL, PP, V-bottom) containing 0.5 mL of LB supplemented with 200 μg/mL kanamycin when necessary. The deep well plates were sealed using a gas-permeable membrane (Diversified Biotech, Breathe-Easy). The following day, a new 96-deep well plate was prepared with 0.45 mL in each well. Then, each well was inoculated to an initial OD_600_ 0.2 by adding 50 μL of OD-normalized pre-culture. The plate was then sealed with a permeable membrane and grown at 30 °C, 200 rpm for 3 h. Each culture was then induced by adding sterile solutions of L-rhamnose, sodium salicylate, or theophylline. To measure uninduced expression, some cultures were left uninduced. The deep well plate was then incubated at 30 °C, 200 rpm for 22 h. Appropriate dilutions were transferred to a 96-well black-walled plate (Greiner, 96-well, PS, F-bottom µCLEAR®). Fluorescence of GFPmut3 was measured from the top at an excitation wavelength of 485 nm and an emission wavelength of 535 nm with a gain of 80 in a BioTek Synergy H1 plate reader. Biomass was determined at 600 nm as scattered light. Each value was blanked using empty media (with or without kanamycin). To determine specific fluorescence, fluorescence intensity was divided by scattered light. To normalize our results, fluorescein sodium salt was used to create a calibration curve. To measure *GFP* expression driven by constitutive promoters, the same protocol was followed without the induction step. A two-sample t-test was used when comparing two samples. When comparing multiple conditions, one-way ANOVA and Tukey’s multiple comparison test were used. ns = not significant, adjusted p-value >0.05; * = adjusted p-value <0.05; ** = adjusted p-value <0.01; *** = adjusted p-value <0.001; **** = adjusted p-value <0.0001. In Figure 3, the results of the statistical analysis are reported in compact letter display: results have statistically different means (adjusted p-value <0.05) if they do not share any letter.

### 2.7. YqjM production in *C. necator*

Pre-cultures were cultivated overnight at 30 °C, 200 rpm in 50 mL tubes containing 5 mL of LB supplemented with 200 μg/mL kanamycin. LB without antibiotics was used to cultivate the wild-type control strains. The following day, 250 mL non-baffled Erlenmeyer flasks were filled with 50 mL of LB supplemented with 200 μg/mL kanamycin. The pre-cultures OD_600_ was measured, and the flasks were inoculated at an initial OD_600_ of 0.1. The cultures were grown at 30 °C, 200 rpm until they reached the induction OD_600_ (0.4, 0.8, 1.2). Then, they were induced by adding L-rhamnose (final concentration: 10 mM). Depending on the experiment, FMN (riboflavin 5′-monophosphate sodium salt hydrate, 73-79%, fluorimetric, Sigma) was added to a final concentration of 1 μM. The cultures were then grown at the chosen temperature (22 °C, 26 °C, 30 °C) for 22 h. The final OD_600_ was measured, then the cultures were centrifuged (3400 x *g*, 30 min, 4 °C), and the wet pellet was weighed. Cell pellets were stored at -20 °C until further analysis. Cell pellets were thawed on ice and resuspended in lysis buffer (20 mM Tris, 150 mM NaCl, pH 7.5, lysozyme), depending on the wet pellet weight (20 mL/g wet pellet). The resuspended cultures were incubated on ice for 20 min, then lysed by sonication (Branson Sonifier 450, intensity 8, duty cycle 40%, 4 cycles of 30 s sonication followed by 1 min on ice). The lysate was then centrifuged (1:15 h, 18 500 x *g*, 4 °C) to obtain the soluble extract.

### 2.8. YqjM production in *E. coli* BL21(DE3)

Pre-cultures were cultivated overnight at 37 °C at 200 rpm in 50 mL tubes containing 5 mL of LB supplemented with 50 μg/mL kanamycin. LB without antibiotics was used to cultivate the control strain. The following day, 250 mL non-baffled Erlenmeyer flasks were filled with 50 mL of LB supplemented with 50 μg/mL kanamycin. Then, the flasks were inoculated at an initial OD_600_ of 0.1. The cultures were grown at 37 °C and 200 rpm until they reached OD_600_ of 0.6. Then, they were induced by adding IPTG (final concentration 1 mM). The cultures were then grown at 20 °C for 20 h. The final OD_600_ was measured, then the cultures were centrifuged (3400 x *g*, 30 min, 4 °C), and the wet pellet weight was measured. Cell pellets were stored at -20 °C until further analysis. Cell pellets were thawed on ice and resuspended in lysis buffer (20 mM Tris, 150 mM NaCl, pH 7.5, lysozyme), depending on the wet pellet weight (20 mL/g wet pellet). The resuspended cultures were incubated on ice for 20 min, then lysed by sonication (Branson Sonifier 450, intensity 8, duty cycle 40%, 4 cycles of 30 s sonication followed by 1 min on ice). The lysate was then centrifuged for 75 min (18 500 x *g*, 4 °C) to obtain the soluble extract.

### 2.9. Determination of FMN concentration

The fresh soluble extract and purified protein samples were used to determine FMN concentration by measuring the absorbance at 452 nm (fresh soluble extract) or 445 nm (purified protein samples) using a spectrophotometer (Jasco V-650, quartz cuvette). To compare the samples, the values were normalized. For each sample, the absorbance volume was multiplied by the buffer volume (to obtain the total FMN produced in the whole culture), then divided by the final OD_600_ (to obtain the specific FMN produced per OD_600_ unit). Soluble extracts from wild-type strains without YqjM expression were used as a blank. A calibration curve using FMN (riboflavin 5′-monophosphate sodium salt hydrate, 73-79%, fluorimetric, Sigma) was used to convert absorbance to FMN concentration values (Figure S10).

### 2.10. SDS-PAGE analysis

Each fresh soluble fraction sample was analysed using sodium dodecyl sulfate-polyacrylamide gel electrophoresis (SDS-PAGE). Samples were prepared by mixing with Laemmli Sample Buffer (Bio-Rad) before incubating at 95 °C for 10 min. Each sample was added to a pre-cast SDS-PAGE gel (Invitrogen, NuPage, 4-12% Bis-Tris, 10 wells). PageRuler (PageRuler™ Prestained Protein Ladder, 10 to 180 kDa, Thermo Scientific™) was used as a protein ladder. The gels were run in fresh MOPS buffer (Invitrogen, NuPAGE MOPS SDS Running Buffer 20x) for 15 min at 50 V, then for 1:10 h at 150 V. The SDS-PAGE gels were stained using Rapid Protein Stain Coomassie Blue (Westburg Life Sciences) and imaged using a ChemiDoc Imaging System (Bio-Rad). To compare the amount of YqjM produced by *C. necator* in each culture condition, the Analyze/Gel tool from Fiji was used to perform gel-based quantification. To minimize the influence of loading variability in the analysis, the intensity of the YqjM band (approximately 40 kDa) was normalized using the intensity of another reference protein band (Figure S6).

### 2.11. Protein purification

Protein purification was performed by using immobilized metal chelate affinity chromatography. Cell pellets of each culture were suspended 10 mL/g wet cell weight (WCW) in washing buffer (sodium phosphate 50 mM, pH 8.0, 300 mM NaCl, 10 mM imidazole). The cells were disrupted by sonication after adding lysozyme and benzonase according to the manufacturer’s instructions and incubation for 30 min in ice (Branson Sonifier 450, intensity 8, duty cycle 40%, 4 cycles of 30 s sonication followed by 1 min on ice). The lysate was then centrifuged for 75 min (18,500 x *g*, 4 °C) to obtain the soluble extract. The cleared supernatant was applied to a 1.5 mL Ni-NTA cartridge (equilibrated on washing buffer) and incubated in ice for 30 min. The column was washed with 10 column volumes (CV) of washing buffer. Finally, the protein was eluted with 5 CV of elution buffer (sodium phosphate 50 mM, pH 8.0, 300 mM NaCl, 250 mM imidazole). Excess imidazole was removed from protein fractions by PD10 desalting columns (Cytiva Europe GmBH, Freiburg im Breisgau, Germany) equilibrated in storage buffer (20 mM potassium phosphate, pH 6.5). The total protein quantity purified from each culture was determined by measuring the absorbance of the pure protein solutions at 280 nm and using the molar extinction coefficient є = 33920 M^-1^ cm^-1^ and the molecular weight of YqjM of 38.41 kDa.

### 2.12. Determination of YqjM activity

Activity of YqjM was measured spectrophotometrically using a BMG Labtech SPECTROstar Omega microplate reader by monitoring NADPH consumption at 340 nm in Sarstedt 96-well plates. Enzyme activity was recorded at 1-minute intervals in 20 mM potassium phosphate buffer (pH 6.5) at 30 °C, using 0.25 mM NADPH and 1 mM 2-cyclohexen-1-one (first dissolved in *i*-PrOH; final concentration 100 mM). Assays were conducted under aerobic conditions. Control reactions containing only molecular oxygen and NADPH were performed, and their corresponding rates were subtracted from those obtained in the presence of 2-cyclohexen-1-one. Specific activities were calculated using ε = 6220 M⁻¹ cm⁻¹ for NADPH and represent the mean of triplicate measurements. All experimental steps, including cultivation, expression, purification, and activity assays, were carried out in biological duplicates for each strain studied.

## 3. Results

### 3.1. T7 RNA polymerase is expressed in *C. necator* H16 under the control of two inducible promoters

*T7RNAP* was integrated into the genome and expressed in *C. necator* under the control of two inducible promoters, induced either by L-rhamnose or salicylate. While the gold standard *E. coli* BL21(DE3) employs an isopropyl β-D-1-thiogalactopyranoside (IPTG)-inducible system, *C. necator* is unable to import lactose or IPTG without expression of a heterologous transporter, making this inducible promoter unsuitable (Gruber et al., 2016). The L-rhamnose-inducible promoter was selected due to its low uninduced gene expression (tightness) and its orthogonal nature in *C. necator* (Alagesan et al., 2018) (yielding strain *C. necator* T7R, Figure 1A and S1). In particular, we selected both the inducible promoter and RBS from the expression plasmid pKRrha, previously used for protein expression in *C. necator*^27^. The native salicylate promoter was selected due to its tightness and high expression range (Hanko et al., 2020) (*C. necator* T7S, Figure 1A and S1).

**Figure 1:**
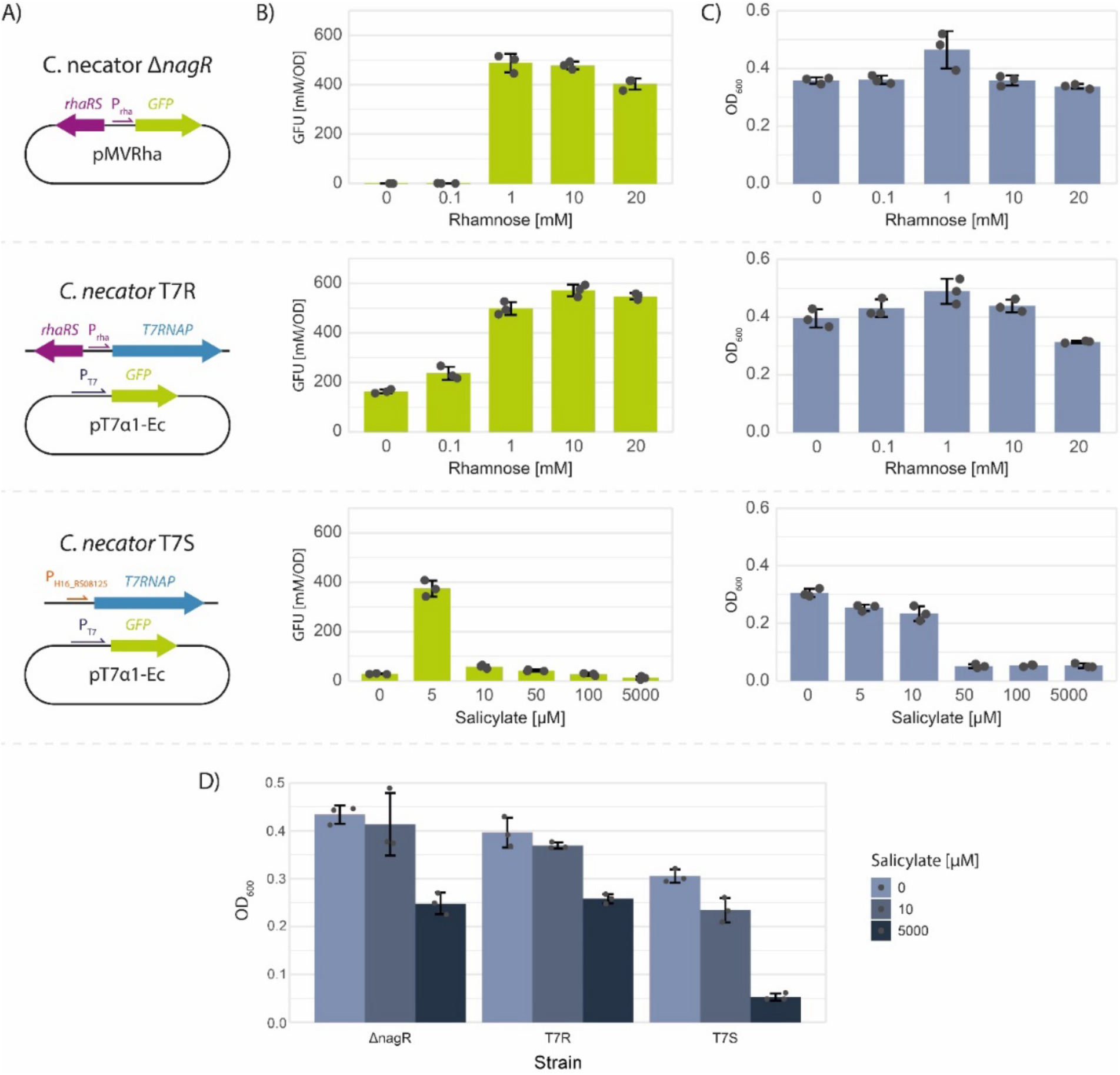
*C. necator* strains containing genomically-integrated T7 RNA polymerase and producing GFP at different levels after induction with either L-rhamnose or salicylate as inducer. A) Genetic constructs introduced by genomic integration and plasmid transformation in the *C. necator* strains. *C. necator* ΔnagR carrying the pMVRha plasmid was used as a benchmark. Diagrams of the genomic context of all strains are reported in Figure S1; B) Specific fluorescence reached after induction with different amounts of inducers, measured by plate reader. C) OD_600_ reached by different *C. necator* strains after addition of different amounts of inducers, measured by plate reader. D) OD_600_ reached by different *C. necator* strains after addition of different amounts of salicylate, measured by plate reader. For all figures, arithmetic means and standard deviations of three biological replicates are reported. Fluorescence is reported as GFU (Green Fluorescent Units: fluorescein equivalent units over OD_600_).

Deletion of the *nagR* gene allows glucose uptake and utilization in *C. necator* (Orita et al., 2012). Thus, it offers an efficient selection method, as the successful deletion-integration is indicated by the glucose-growing phenotype. We decided to integrate the two genetic constructs into the *nagR* locus, while deleting the *nagR* gene. The *nagR* gene was also deleted from the parental strain *C. necator* ΔRM resulting in the strain *C. necator ΔnagR*, used as a negative control.

After integration of the two constructs, GFPmut3 (GFP) fluorescence was used as a quantitative readout to determine protein expression level. Initially, *GFP* expression was driven by a plasmid under the control of a T7 consensus promoter and RBS B0034m (RBS 1), previously investigated in *C. necator* (Keating and Young, 2023). We selected the *GFP* genetic sequence from the GoldenStandard public library (GFP (Ec)) (Blázquez et al., 2022). We also expressed the same gene using our recently developed plasmid pMVRha as a benchmark (Vajente et al., 2024) (Figure 1A).

GFP fluorescence was successfully detected in both T7 strains after induction (Figure 1B). Using concentrations of salicylate higher than 10 μM impaired growth in the strain *C. necator* T7S, leading to lower final OD_600_ (Figure 1C). High concentrations of salicylate decreased final OD_600_ in the other strains as well, but not as drastically (Figure 1D). This suggests that a feedback mechanism may be responsible for the growth inhibition. However, this behavior was not observed in previous studies with the same promoter and may therefore be attributed to the expression of T7RNAP using a native promoter (Hanko et al., 2020). We decided not to pursue this strain further due to the potential impact of the native metabolism in future experiments.

The L-rhamnose-inducible promoter yielded the highest *GFP* expression at 10 mM inducer concentration. However, the maximum fluorescence was comparable to that of our previously developed pMVRha expression plasmid. This is intriguing, since T7RNAP usually delivers high mRNA levels of the gene of interest (Studier and Moffatt, 1986). We speculated that the high level of mRNA produced by T7RNAP was not successfully translated into protein, creating a bottleneck in protein production. Thus, we explored the potential barriers in mRNA translation.

### 3.2. *C. necator* contains more rare codons compared to *E. coli*

We speculated that mRNA translation could be limited by the cytoplasmic pool of charged tRNAs (non-optimal codon usage). To investigate this hypothesis, we designed a GFP genetic sequence codon-optimized for *C. necator* (GFP(Cn)). While desirable, codon harmonization of this gene was impossible, as that required the source organism genome which was unavailable at the time. We then calculated the Codon Adaptation Index (CAI) of this new codon-optimized sequence for both *E. coli* and *C. necator* (Sharp and Li, 1987). Surprisingly, the *C. necator* codon-optimized sequence also showed high CAI for *E. coli* (Figure 2A). Thus, sequences that were optimal for *C. necator* were also optimal for *E. coli*, but not *vice versa*. This discrepancy could be traced to the codon usage of both organisms. We observed that for each amino acid, in *E. coli* the synonymous codons were used in a balanced manner. In contrast, it was common to encounter amino acids with imbalanced codon distributions in *C. necator*, with one codon prevalent and the others barely present in the genome. An example of this codon use is shown in Figure 2B (the complete analysis can be found in Table S4). To quantify this phenomenon, the frequency of each codon was reported in a histogram (Figure 2C). In *C. necator*, 23 codons are rare (frequency lower than 10%), whereas in *E. coli*, this number is six. The high number of rare codons suggests that many charged tRNAs may be present in lower quantities in *C. necator*, creating potential bottlenecks during protein production.

**Figure 2:**
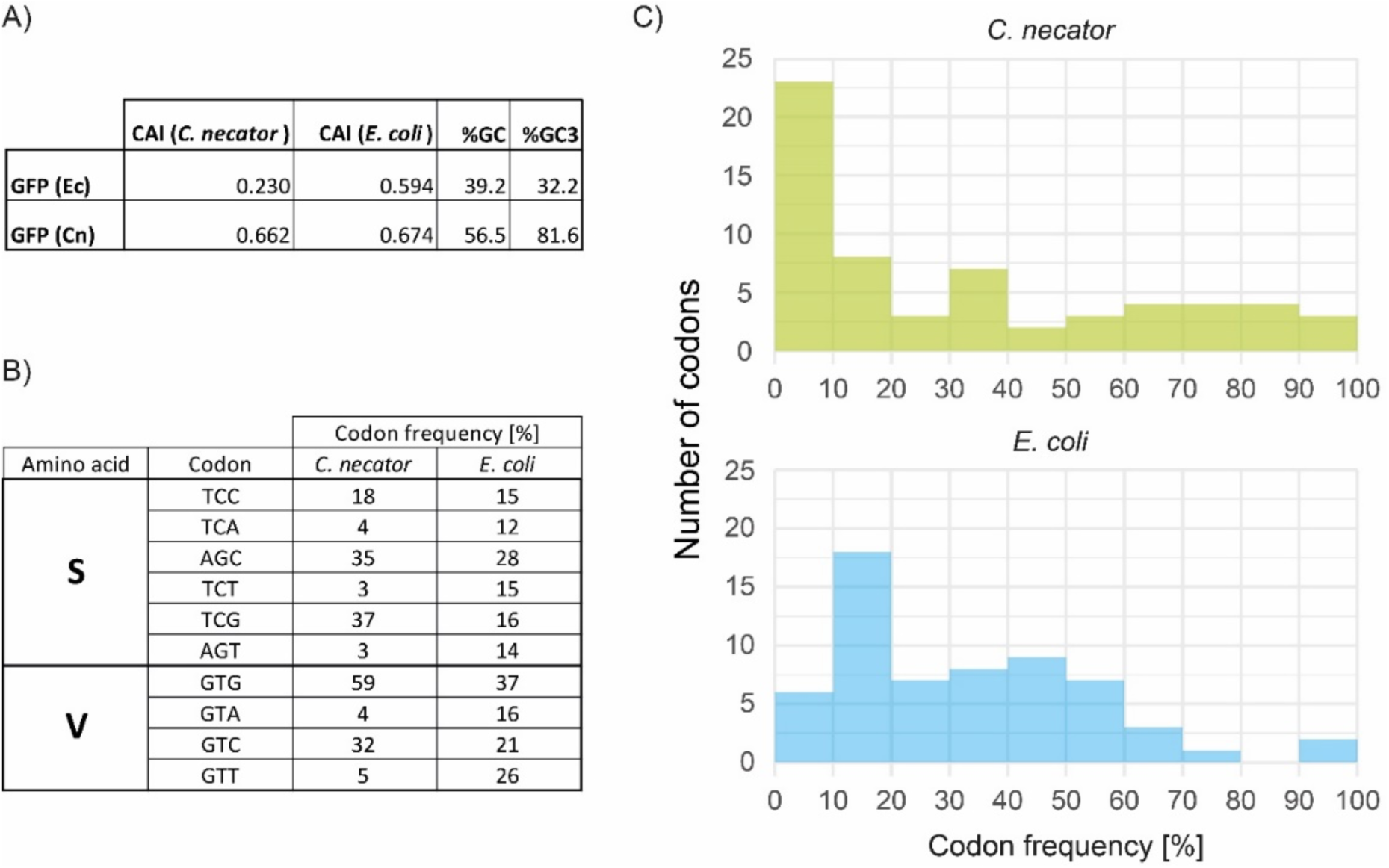
*C. necator* contains a higher number of rare codons compared to *E. coli*. A) Codon Adaptation Index (CAI) and GC percentage (both total - %GC; and of the third base of the codon - %GC3) of the two GFP genetic sequences in *E. coli* and *C. necator*. The original GFP sequence (GFP (Ec)) showed high CAI for *E. coli*, while the sequence optimized for *C. necator* (GFP (Cn)) showed high CAI for both organisms. B) Codon frequency of synonymous codons encoding serine and valine in both *E. coli* and *C. necator*. C) Number of codons present at each frequency in *C. necator* and *E. coli*. In *E. coli*, most codons are in the intermediate range, showing a balanced distribution. In *C. necator*, more codons are present at low and high frequencies.

### 3.3. Codon usage is the limiting factor for high-yield protein production in *C. necator*

To confirm whether the mRNA translation was limited by low ribosome occupancy (RBS strength) or charged tRNA depletion (codon usage), we assembled multiple genetic constructs and quantified GFP production. First, we selected two RBSs previously investigated in *C. necator*: RBS B0034m (used in Figure 1) (RBS 1) and RBS T7g10, previously used to drive gene expression in the plasmid pKRrha (RBS 2) (Figure 3A)^27, 30^. We determined their relative strength in *C. necator* by measuring GFP production under the control of the constitutive promoter J23106 (Figure 3B), and observed that RBS 2 was stronger than RBS 1 under constitutive expression in our parental strain *C. necator* ΔRM. We then used these two RBSs to drive GFP(Ec) expression in our *C. necator* T7R strain under the control of the T7 consensus promoter (T7α) (Figure 3C). Increasing the RBS strength only slightly increased final fluorescence. Using the stronger RBS also increased uninduced *GFP* expression (Figure 3C, plasmids pT7α1/2-Ec). We then evaluated the influence of codon tuning towards protein production by expressing two different GFP genetic sequences encoding the same protein: the original sequence from the GoldenStandard library (GFP(Ec)) and the codon-optimized sequence (GFP(Cn)). We could observe a striking increase in fluorescence (4-fold) when the codon-optimized GFP(Cn) gene was used, showing that mRNA translation was probably impaired by tRNA depletion (Figure 3C, plasmids pT7α1-Ec/Cn), as previously hypothesized. Even when cloning the two sequences in the pMVRha expression plasmid, codon optimization resulted in a 2-fold increase in final fluorescence. Therefore, tRNA depletion was a bottleneck in protein production in both our T7-driven expression system as well as our normal L-rhamnose-driven expression plasmid.

**Figure 3:**
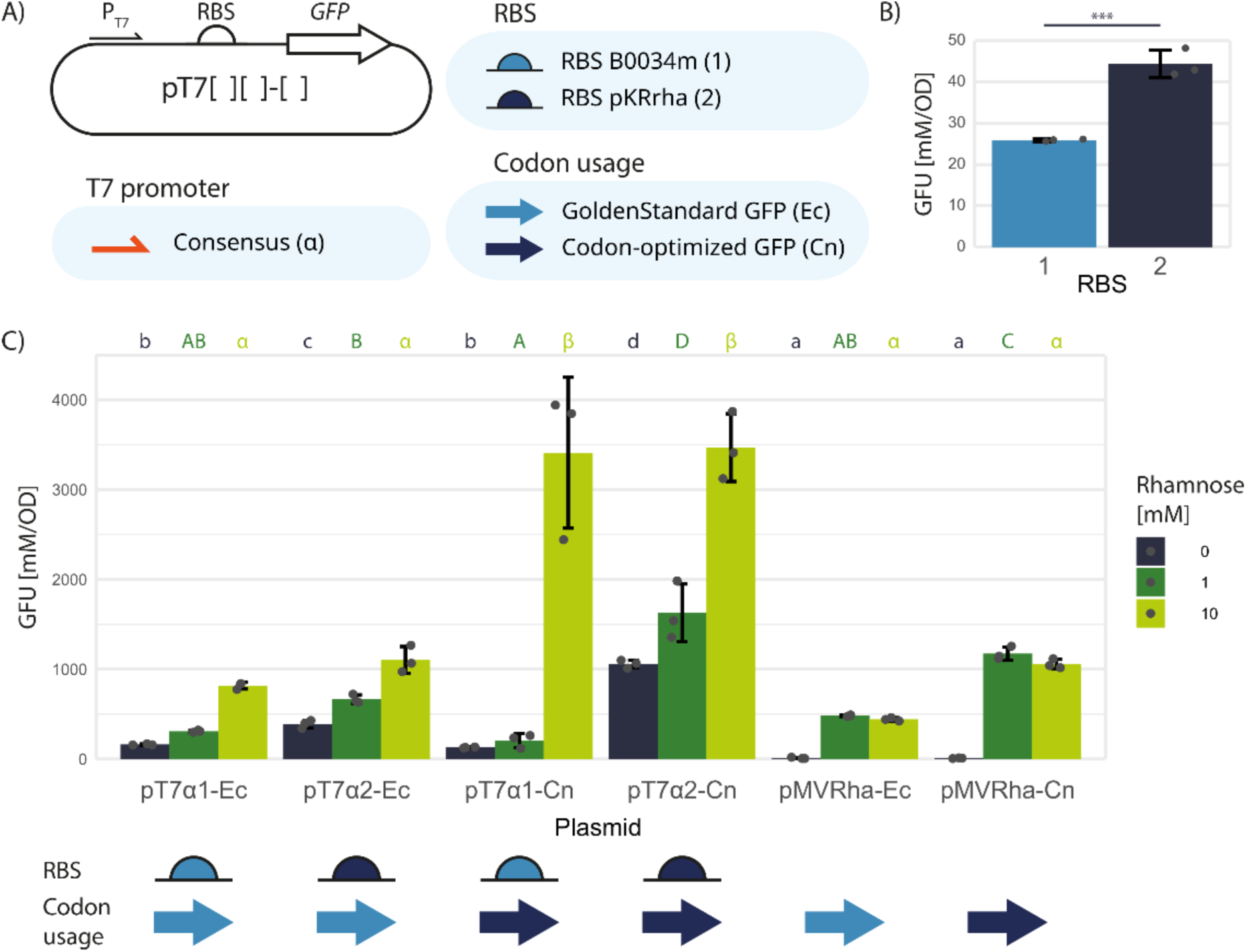
GFP protein production in *C. necator* increases substantially after codon optimization. A) Genetic parts analyzed and schematic representation of the test plasmid used. B) Specific fluorescence reached by *C. necator* ΔRM expressing the codon-optimized *GFP* gene under the control of the constitutive promoter J23106 and two different RBSs (B0034m, pKRrha); C) Specific fluorescence reached by *C. necator* T7R expressing the codon-optimized *GFP* gene (*GFP* (Cn)) or the non-optimized sequence from the GoldenStandard library (*GFP* (Ec)) under the control of promoter T7α and different RBSs. Arithmetic means and standard deviations of three biological replicates are reported. Fluorescence is reported as GFU (Green Fluorescent Units: fluorescein equivalent units over OD_600_).

### 3.4. Newly designed T7 expression strains reduced leaky expression of the system

We successfully improved the maximum GFP production in our *C. necator* T7R strain by using an expression plasmid containing a codon-optimized *GFP* gene, a consensus T7 promoter, and a strong RBS (pT7α2-Cn). This combination achieved a 3.5-fold improvement in fluorescence compared to our expression plasmid pMVRha (Figure 3C). However, the uninduced expression of the T7 system was higher than that of pMVRha. We thus proceeded to optimize the integrated *T7RNAP* genetic construct to reduce leaky expression. Our original system contained a non-codon-optimized *T7RNAP* sequence under the control of a strong RBS (RBS 2). We hypothesized that codon-harmonizing the gene and selecting a weaker RBS could influence both induction strength and tightness, and therefore a new strain with these modifications (*C. necator* T7R2, Figures 4A and S1) was constructed. In addition, we designed a new genetic circuit using a Self-splicing Intron-Based Riboswitch (SIBR). This element was recently employed to tightly regulate endonuclease expression in *C. necator* (Della Valle et al., 2025). Following that example, we introduced the SIBR inside the *T7RNAP* coding sequence to decrease uninduced expression of *T7RNAP*. When theophylline is added to the medium, the SIBR undergoes a self-catalyzed excision, resulting in a reconstitution of the complete T7RNAP coding sequence.

**Figure 4:**
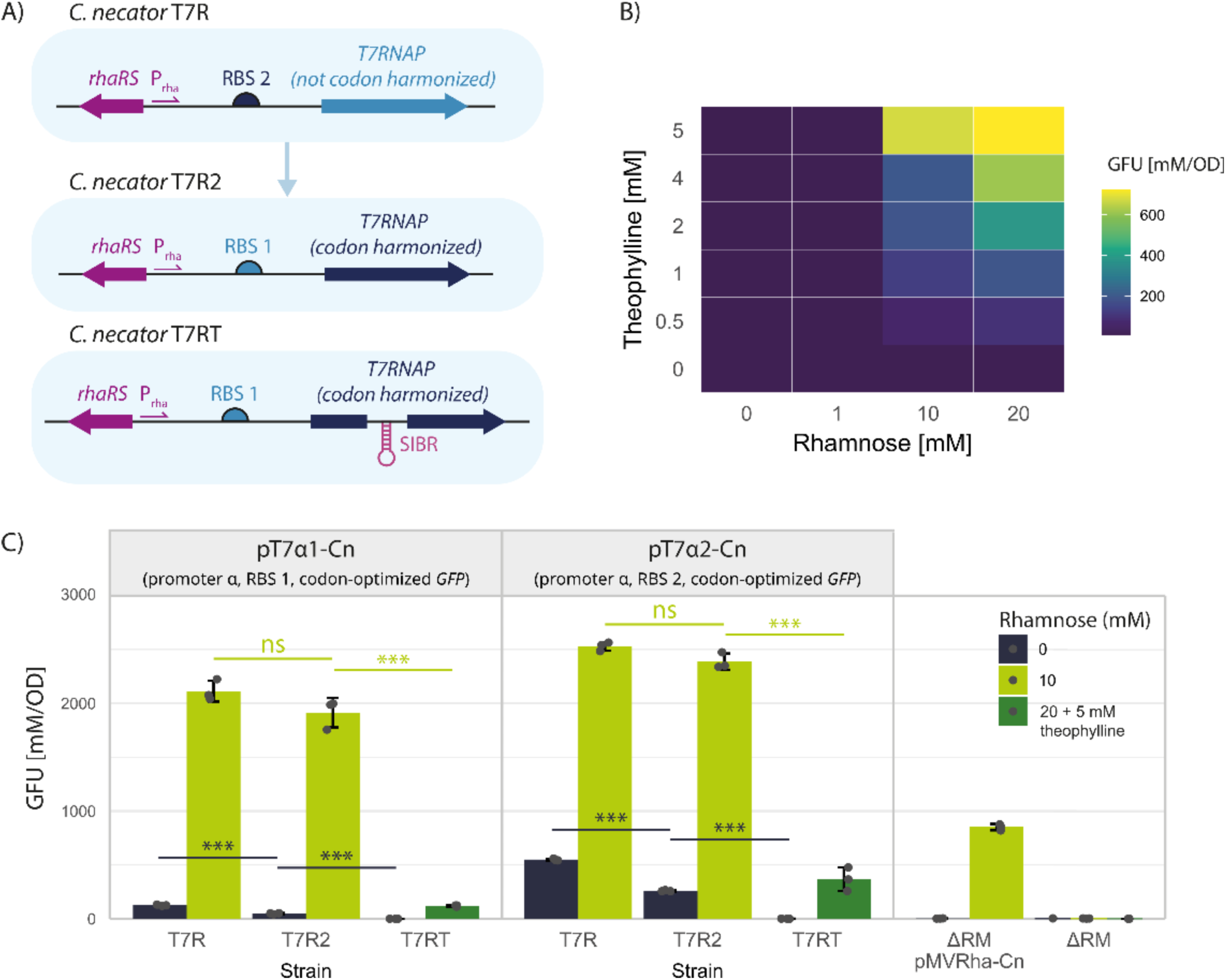
*C. necator* T7R2 maintains maximum GFP production while showing lower uninduced fluorescence compared to the parent strain. A) Genetic circuits introduced by genomic integration in the new *C. necator* strains. Schematic representations of the genomic context of all strains are reported in Figure S1; B) Specific fluorescence reached by strain *C. necator* T7RT carrying plasmid pT7α2-Cn (T7 consensus promoter, RBS 2, codon-optimized *GFP*) after induction with different amounts of L-rhamnose and theophylline; C) Specific fluorescence reached by different *C. necator* strains carrying plasmids pT7α1-Cn, pT7α2A-Cn, or pMVRha-Cn (codon-optimized *GFP*, control). Arithmetic means and standard deviations of three biological replicates are reported. Fluorescence is reported as GFU (Green Fluorescent Units: fluorescein equivalent units over OD_600_).

We designed and assembled a strain with *T7RNAP* under both L-rhamnose and SIBR regulation (*C. necator* T7RT, Figures 4A and S1). Both novel genetic circuits were integrated near the *nagR* locus. Compared to our initial strains, we chose not to delete *nagR* in our final strains, to remove potential metabolic burden or other influences caused by *nag* operon de-repression. We then transformed *C. necator* T7RT with the best-performing plasmid pT7α2-Cn and induced *GFP* expression with different amounts of L- rhamnose and theophylline to identify the optimal inducer concentrations (Figure 4B). *GFP* expression proved to be tight, requiring high concentrations of both inducers to observe fluorescence production. This is consistent with the reported data from a similar system in *E. coli* (Della Valle et al., 2025). However, the maximum fluorescence obtained with strain *C. necator* T7RT was drastically lower than the one obtained with the other strains. Conversely, strain *C. necator* T7R2 maintained similar maximum fluorescence to *C. necator* T7R while reducing uninduced expression by half (Figure 4C).

### 3.5. Optimization of genetic parts in the expression plasmid can further improve the system

After selecting *C. necator* T7R2 as chassis, we focused on the optimization of the genetic parts in the expression plasmid. We assessed two new T7 promoter variants identified in previous studies for their higher performance compared to the consensus promoter (Figure 5A) (Conrad et al., 2020; Jones et al., 2015). Both new T7 promoters were able to maintain the maximum protein production; one variant also decreased uninduced expression (Figure 5B). In addition, we designed a synthetic RBS derived from the highly efficient RBS of the pET plasmid series, changing the Epsilon sequence to reflect the rRNA sequence in *C. necator* (RBS 3) (Figure S5) (Olins and Rangwala, 1989). However, this RBS did not increase *GFP* expression compared to RBS 1 under the control of a constitutive promoter (Figure 5C). When we expressed *GFP* (Cn) in our strain *C. necator* T7R2 under the control of the consensus T7 promoter and RBS 3, the maximum fluorescence did not change drastically, while uninduced expression almost doubled compared to the strong RBS 2 (Figure 5D). These results confirmed our previous findings that increasing RBS strength increases system leakiness. Increasing RBS strength from RBS 1 to RBS 2 increased both uninduced and induced expression. However, when we increased the RBS strength further, from RBS 2 to RBS 3, only the uninduced expression increased significantly. This could indicate that another bottleneck in protein production has been reached. Optimizing both the integrated genetic circuit and the plasmid-based expression was thus essential to achieve a system with high maximum and low uninduced expression.

**Figure 5:**
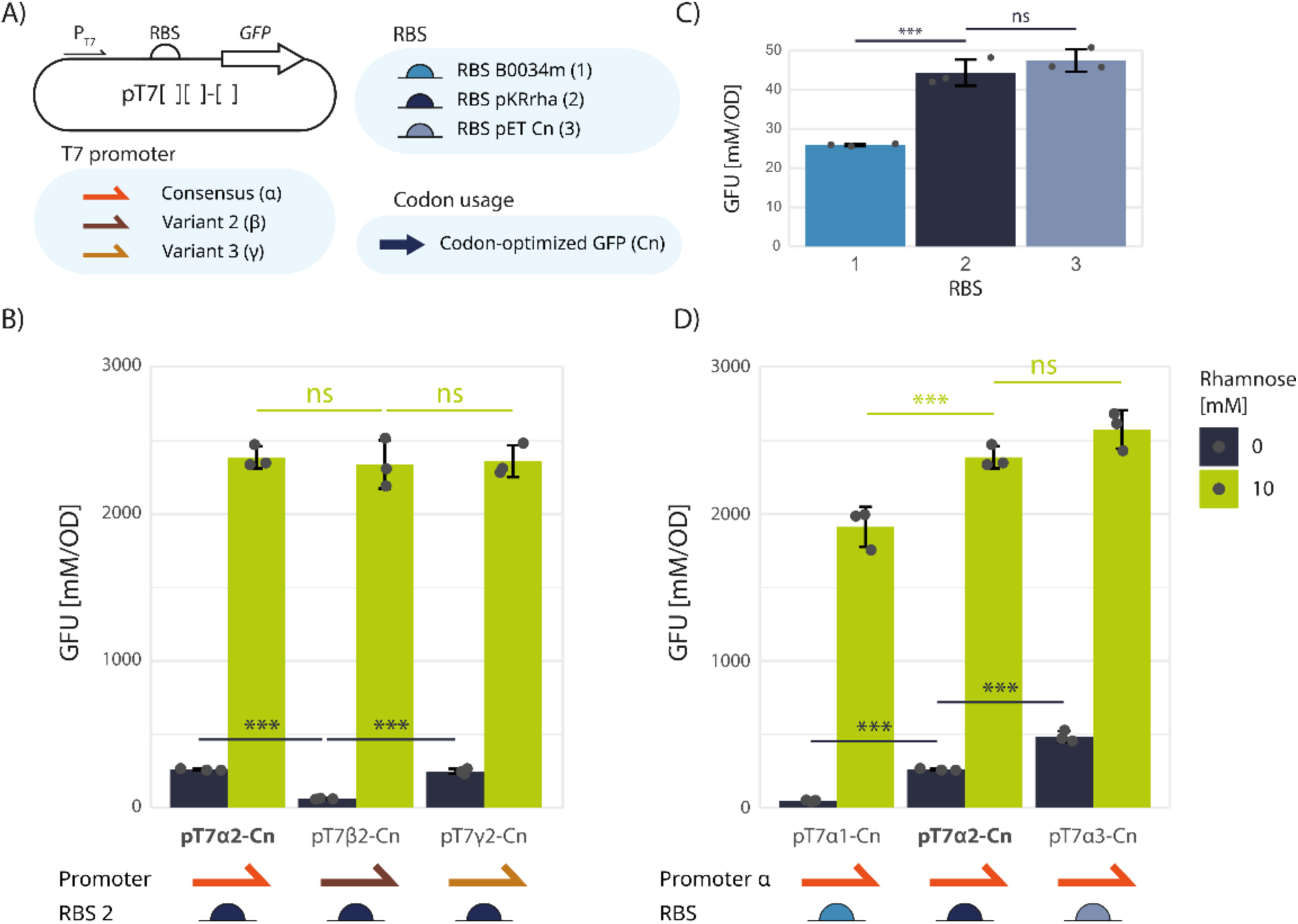
An optimized T7 promoter decreased leakiness while maintaining maximum fluorescence production. A) Genetic parts analyzed and schematic representation of the test plasmid used. B) Specific fluorescence reached by *C. necator* T7R2 expressing *GFP* (Cn) under the control of different T7 promoter variants and RBS 2. Plasmid pT7α2-Cn (bold) is used as a reference. C) Specific fluorescence reached by *C. necator* ΔRM expressing *GFP* (Cn) under the control of constitutive promoter J23106 and the three different RBSs. D) Specific fluorescence reached by *C. necator* T7R2 expressing *GFP* under the control of promoter T7α and three different RBSs. Plasmid pT7α2-Cn (bold) is used as reference. Arithmetic means and standard deviations of three biological replicates are reported. Fluorescence is reported as GFU (Green Fluorescent Units: fluorescein equivalent units over OD_600_).

### 3.6. Ene-reductase YqjM can be produced at high titer in *C. necator*

After producing GFP as a model protein with an easy-to-follow readout, we used our optimized strain for enzyme production. We selected YqjM, a FMN-dependent ene-reductase from *Bacillus subtilis*, as a target enzyme due to its previous production in both *E. coli* (Pesic et al., 2017) and *C. necator* (Assil-Companioni et al., 2019). We used our optimized system (*C. necator* T7R2, carrying plasmid pT7α2 with *yqjM* codon-harmonized for *C. necator*) to produce YqjM. We also assessed the most favorable culture conditions for enzyme production in *C. necator*. Thus, cultures were induced at different ODs (0.4, 0.8, 1.2) and incubated at various temperatures after induction (22 °C, 26 °C, 30 °C). In addition, we used our parental strain *C. necator* ΔRM carrying the pMVRha expression plasmid as a benchmark. We expressed both *E. coli* codon-optimized (Ec) and *C. necator* codon-harmonized (Cn) *yqjM* genes in pMVRha to confirm whether codon usage was as relevant to enzyme production as it was to GFP production. To compare our systems with a widely used strain, we produced the same protein in *E. coli* BL21(DE3) carrying the *E. coli* codon-optimized gene in plasmid pET28a.

YqjM forms a tetramer in its active form, with one FMN molecule binding to each subunit (Kitzing et al., 2005), and only the FMN-loaded fraction is predicted to be catalytically active (Theorell et al., 1957). It has previously been shown that recombinant YqjM produced in *E. coli* is not fully saturated with FMN (Fitzpatrick et al., 2004, 2003). Consequently, in biocatalytic applications, purified YqjM (Pesic et al., 2017) and related ene-reductases (Tonoli et al., 2023) are often incubated with free FMN prior to use to restore full activity through cofactor reconstitution. Therefore, we quantified both FMN (Figure 6A) and soluble protein (Figure 6B, Figure S6) produced by the microbial cultures. All the strains were able to produce soluble YqjM and FMN, albeit at different levels. *E. coli* BL21(DE3) achieved the highest production of soluble protein (Figure 6B). However, *C. necator* produced higher amounts of FMN compared to *E. coli* BL21(DE3) (Figure 6A) from the same culture volume. The highest amount of FMN (total FMN) was produced with our previously developed pMVRha plasmid carrying the codon-harmonized *yqjM* gene due to its high final OD_600_ (Figure S7). Furthermore, using the codon-harmonized gene increased both FMN and soluble protein production compared to the non-codon-adjusted gene, confirming the importance of codon usage tuning. Our optimized strain *C. necator* T7R2 reached low final OD_600_; however, the FMN quantity produced per OD_600_ unit (specific FMN) was the highest among the tested strains, indicating a higher concentration of FMN cofactor in the microbial biomass (Figure 6A).

**Figure 6:**
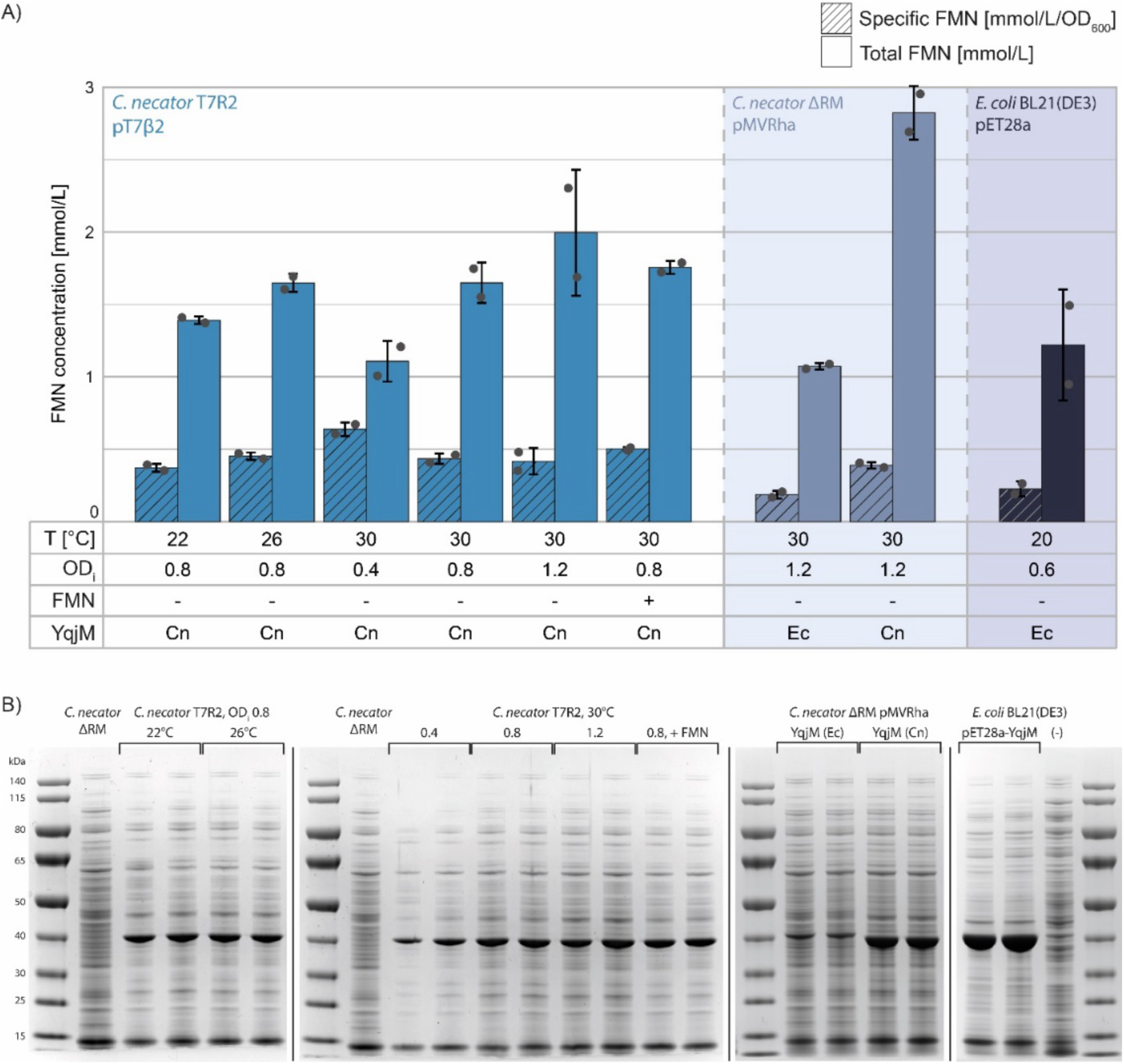
The optimized *C. necator* T7R2 strain produces soluble and FMN-loaded YqjM. A) FMN concentration measured in the soluble extract of *C. necator* T7R2, *C. necator* ΔRM, and *E. coli* BL21(DE3) under different expression conditions. To calculate the specific FMN production, the total FMN quantity was divided by the final OD_600_ reached by each culture. FMN absorbance of each wild-type strain without expression plasmid was used to blank the values. T[°C]: incubation temperature after induction; OD_i_: induction OD; FMN: in one experiment (+), FMN was supplemented to the culture at a final concentration of 1 μM; YqjM: two sequences with different codon usages were used (Ec, Cn) (see Table S5 for CAI and Table S3 for sequences). Arithmetic means and standard deviations of two biological replicates are reported. FMN quantity is reported in mmol_FMN_/L of culture. B) SDS-PAGE analysis of the soluble extracts analyzed in Figure 6A. A gel-based quantification of soluble protein production is reported in the Supplementary Information (Figure S6).

### 3.7. *C. necator* is more efficient in cofactor loading of YqjM

Since *C. necator* was able to produce higher amounts of intracellular FMN, we wondered whether this FMN was successfully incorporated into the heterologous enzyme, yielding FMN-loaded YqjM. Proteins produced by different strains under optimal conditions were isolated and evaluated for catalytic activity. These recombinant enzymes were purified and analyzed to (1) determine protein yield per culture volume, (2) quantify FMN content, and (3) assess catalytic activity without FMN supplementation, thereby reflecting the intrinsic efficiency of each host system.

His-tagged recombinant YqjM was purified from the soluble fraction of cell lysates using Ni-NTA agarose, followed by desalting with PD-10 columns. The protein was obtained in good yields (158 mg/L, 47 mg/L, and 148 mg/L for *E. coli*, *C. necator* T7R2, and *C. necator* ΔRM cultures, respectively; or 8.3 mg/OD_600_, 5.8 mg/OD_600_, and 4.6 mg/OD_600_, Table 1). Although *E. coli* produced higher total protein levels, the extent of FMN incorporation differed substantially. FMN saturation was assessed spectrophotometrically by measuring absorbance at 445 nm in purified protein solutions. Significantly higher FMN concentrations per mg of protein were observed for enzymes expressed in *C. necator* T7R2 (16.7 nmol/mg) and *C. necator* ΔRM (11.5 nmol/mg), whereas *E. coli*-derived protein showed considerably lower values (3.2 nmol/mg), indicating reduced cofactor loading. To demonstrate enhanced FMN incorporation, enzyme activity was evaluated using an NADPH-dependent cyclohexenone reduction assay (Figures S8, S9). Enzymes expressed in *C. necator* exhibited 1.6- and 1.4-fold higher activity compared to the *E. coli*-derived YqjM, indicating that improved flavin loading translates directly into enhanced catalytic efficiency.

**Table 1.**
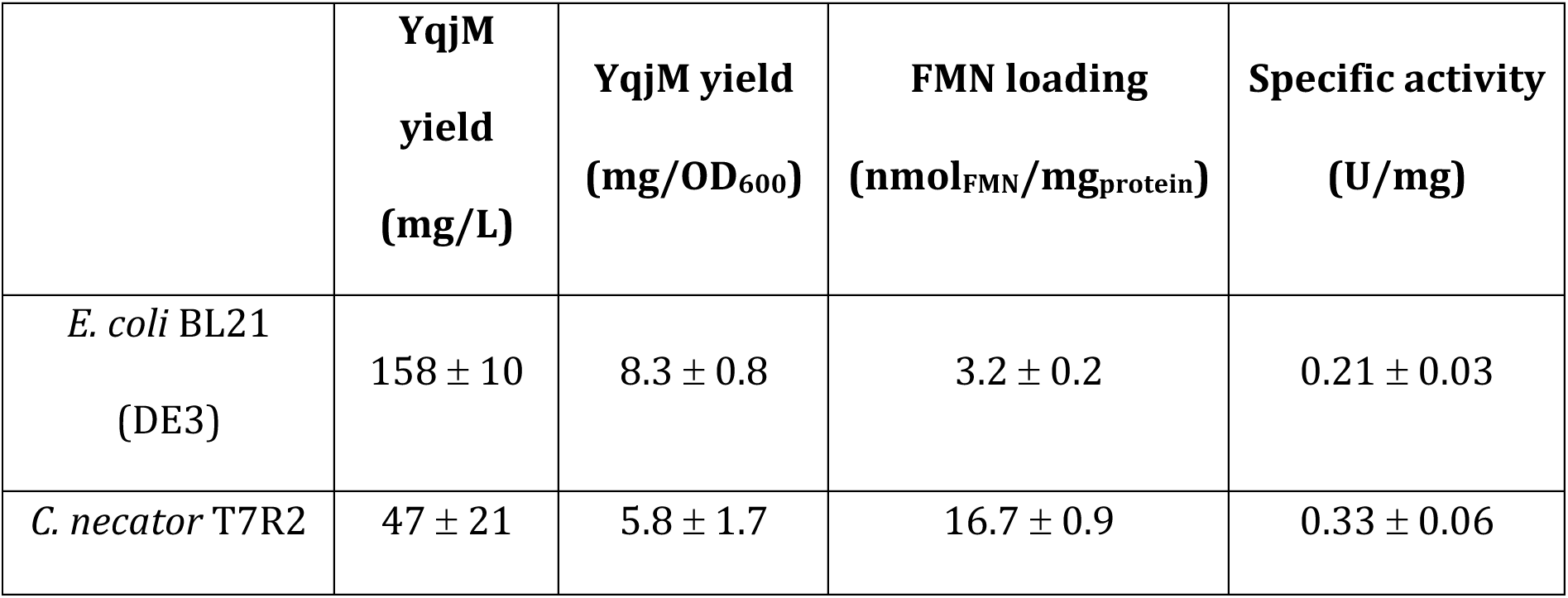
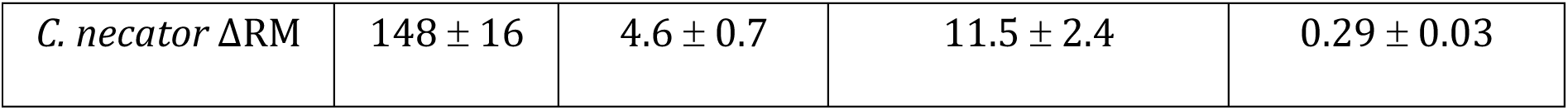
The results of purification of YqjM and the corresponding activities of biological duplicates. The total protein quantity (YqjM) for each culture was determined by measuring the absorbance of the pure protein solutions at 280 nm and using the molar extinction coefficient ε of 33920 M^-1^ cm^-1^ and the molecular weight (MW) of 38.41 kDa. The protein quantity obtained per OD_600_ was divided by the final OD_600_ reached by each culture. To calculate the specific FMN production, the total FMN quantity was divided by the final OD_600_ reached by each culture. FMN absorbance of each wild type strain without expression plasmid was used to blank the values. Specific activities were then recorded spectrophotometrically from the decrease of NADPH in YqjM catalyzed reduction of 2-cyclohexen-1-one. Reaction conditions: 30 °C; 20 mM potassium phosphate buffer (pH 6.5); 0.25 mM NADPH; 0.5 μM YqjM, 1 mM 2-cyclohexen-1-one in *i*-PrOH (100 mM final concentration). Specific activities were calculated from the mean values of triplicate measurements by subtracting the activity measured in the presence of the substrate and the activity measured with molecular oxygen as only substrate.

|  | **YqjM**<br>**yield**<br>**(mg/L)** | **YqjM yield**<br>**(mg/OD<sub>600</sub>)** | **FMN loading**<br>**(nmol<sub>FMN</sub>/mg<sub>protein</sub>)** | **Specific activity**<br>**(U/mg)** |
| --- | --- | --- | --- | --- |
| *E. coli* BL21<br>(DE3) | 158 ± 10 | 8.3 ± 0.8 | 3.2 ± 0.2 | 0.21 ± 0.03 |
| *C. necator* T7R2 | 47 ± 21 | 5.8 ± 1.7 | 16.7 ± 0.9 | 0.33 ± 0.06 |
| *C. necator* ΔRM | $148 \pm 16$ | $4.6 \pm 0.7$ | $11.5 \pm 2.4$ | $0.29 \pm 0.03$ |

## 4. Discussion

Given the high potential of *C. necator* for biocatalytic applications and C1-based manufacturing, there is an urgent need to develop more efficient protein production systems for this non-model host. In this study, we successfully integrated T7RNAP driven by two different inducible promoters. The native salicylate-based inducible promoter (NahR) was induced by micromolar amounts of inducer; however, higher concentrations inhibited cell growth. Notably, this phenomenon is not due to salicylate toxicity, as previous studies have shown that salicylate can have a beneficial growth effect of up to 5 mM (Hanko et al., 2020). However, this was not the case in our study, even for the non-engineered strains. Growth inhibition was unlikely to be due to the metabolic burden of T7RNAP or GFP production, since the cultures with impaired growth did not exhibit increased GFP fluorescence (Figure 1).

Another attempt to use a heterologous salicylate promoter in *C. necator* achieved no protein production, further demonstrating that different genetic systems may interact differently with the native metabolism (Mishra et al., 2024). As our aim was to create a robust and reliable system, we opted to abandon the salicylate promoter and focused on the L-rhamnose-inducible strain instead. While cell growth remained consistent across all inducer concentrations, the maximum fluorescence achieved with the T7-based system was similar to that obtained with plasmid pMVRha.

We speculated that the high level of mRNA produced by T7RNAP was not being successfully translated into protein, creating a bottleneck in protein production. Thus, we focused our investigation on two variables: RBS strength and codon usage of the gene of interest. Using a stronger RBS increased uninduced expression considerably in our system while only slightly increasing maximum GFP fluorescence (Figures 3 and 5). This trade-off was disadvantageous, because regulatory tightness is an essential quality in a production strain. Typically, separating the biomass accumulation phase from the protein production phase reduces the metabolic burden during growth, increases plasmid retention and final biomass, and leads to a higher protein yield(De Baets et al., 2024).

While the role of codon usage in heterologous protein production has proven elusive in *E. coli*, with different proteins benefiting from codon optimization, harmonization, or non-optimized sequences(Claassens et al., 2017), the importance of this variable has been emphasized repeatedly(Liu et al., 2021). We found that most amino acids in *C. necator* show a strong bias towards specific codons. Conversely, *E. coli*’s synonymous codons are distributed more evenly (Figure 2). Each rare codon (used less than 10%) can impede protein production, and *C. necator* has almost four times more rare codons than *E. coli* (Figure 2). The main bottleneck in protein production was indeed caused by the gene of interest’s suboptimal codon usage. Optimizing the sequence of *GFP* increased the maximum fluorescence by 4-fold compared to the non-optimized gene (Figure 3). Interestingly, our plasmid control also benefited from codon optimization, with a 2-fold increase in fluorescence. This indicates that protein translation was already a bottleneck when using the pMVRha plasmid. Another recent study also observed the important role of codon tuning in protein production in *C. necator* (Mishra et al., 2024), and we confirmed that both codon optimization (GFP, Figure 3) and codon harmonization (YqjM, Figure 6) improved protein levels considerably. By simply tuning the codon usage of the YqjM genetic sequence, the production of soluble enzyme using the pMVRha plasmid increased by more than 3-fold (Figure 6B and S6).

Building on this knowledge, we developed two new T7 strains carrying a codon-harmonized *T7RNAP* gene under the control of a weaker RBS. *C. necator* T7R2 showed a 50% reduction in uninduced GFP production (Figure 4). However, an attempt to increase tightness further by using a newly developed theophylline-responsive system (SIBR2.0)(Della Valle et al., 2025) was unsuccessful because the system was tight but not responsive to induction. This was unexpected because T7RNAP only requires low expression to drive high-level mRNA transcription.

To further optimize our expression plasmid, we selected a T7 promoter variant that reduced uninduced expression and generated a tighter system (Figure 4). Our in-depth investigation of the genetic elements involved in the system revealed the importance of each variable’s role and helped to identify the bottlenecks limiting the desired phenotype.

Ultimately, we challenged our strain with the production of the flavin-dependent ene-reductase YqjM from *Bacillus subtilis* and compared it with the gold standard *E. coli* BL21(DE3) pET system. Our optimized T7-based system (*C. necator* T7R2 pT7β2) successfully produced soluble YqjM (Figure 6B) and achieved the highest production of FMN per biomass unit (Figure 6A), resulting in a high intracellular amount of cofactor-loaded YqjM (Table 1). However, T7-based expression led to a considerable decrease in final cell density compared to *C. necator* ΔRM carrying the expression plasmid pMVRha (Figure S7). This T7RNAP-induced growth impairment was not present when producing GFP, probably indicating an increased metabolic burden during the production of more complex proteins. Optimizing the media composition and feeding strategy may ameliorate the growth burden of *C. necator* T7R2. When producing YqjM using the pMVRha plasmid, the final yields of both soluble protein and FMN were higher than those obtained with our T7-based strains due to the higher final OD_600_. A similar result was previously observed in *E. coli*, where some expression plasmids outperformed commonly used T7-based expression systems (Schuster and Reisch, 2022). *E. coli* BL21(DE3) carrying the pET28a plasmid achieved the highest titers of soluble YqjM (Table 1). However, the cofactor production was lower than in *C. necator*, resulting in a lower fraction of active protein (Figure 6A, Table 1). YqjM purified from *C. necator* showed higher FMN-loading (3- and 5-fold increase) and specific activity (1.4- and 1.6-fold increase) compared to YqjM purified from *E. coli*. These results demonstrate that selecting an alternative host can inherently overcome cofactor incorporation limitations. This is particularly advantageous for biocatalytic and fermentation-based processes, where suboptimal cofactor loading can compromise overall productivity. While partial activity loss in *E. coli*-expressed enzymes can be mitigated through post-purification FMN supplementation (Classen et al., 2014; Pesic et al., 2017), that is unfeasible in whole-cell applications. Thus, host optimization represents a powerful strategy to improve the performance of flavin-dependent biocatalysts without requiring additional processing steps.

In conclusion, our study demonstrates that *C. necator* is a promising industrial chassis for protein production. The T7-based system developed herein resulted in higher FMN-bound YqjM production per unit of biomass, whereas our previously developed expression plasmid pMVRha resulted in higher total FMN-bound YqjM production due to higher cell density achieved at the end of the growth phase. These systems could be used to produce other enzymes that are difficult to express in *E. coli* due to the formation of inclusion bodies (Srinivasan et al., 2002) or a lack of cofactor maturation chaperones (Ryu et al., 2024). While *E. coli* BL21 produced higher amounts of soluble enzyme, our system outperformed *E. coli* in expressing FMN-bound enzyme, further highlighting the potential of *C. necator* for biocatalysis applications. Furthermore, these optimized expression systems could be used for lithoautotrophic or formatotrophic enzyme production, ultimately leading to CO_2_-driven enzyme synthesis.

## Declaration of competing interest

The authors declare that they have no known competing financial interests or personal relationships that could have appeared to influence the work reported in this paper.

## Acknowledgements

We thank Riccardo Clerici, Franziska Meier and all the employees at Formo GmbH for support and discussions. This project has received funding from the European Union’s Horizon 2020 research and innovation programme under the Marie Skłodowska Curie grant agreement No. 955740 and the Netherlands Organization for Scientific Research (OCENW.XS23.3.002).

## Author contributions

### CRediT

Matteo Vajente (RUG) – Conceptualization, Investigation, Writing – original draft Marleen Hallamaa (RUG) – Investigation, Writing – original draft Hendrik Ballerstedt (RWTH-Aachen) – Supervision, Writing – review & editing Lars M. Blank (RWTH-Aachen) – Resources, Supervision, Writing – review & editing Sandy Schmidt (RUG) – Conceptualization, Supervision, Resources, Writing – review & editing

### Funding sources

This work was supported by the European Union’s Horizon 2020 research and innovation programme under the Marie Skłodowska Curie grant agreement No. 955740 and the Netherlands Organization for Scientific Research (OCENW.XS23.3.002).

## Appendix A. Supplementary data

The supplementary material provides supporting data and figures, further details on the strains and plasmids used, sequences, and supporting data (e.g. codon Adaptation Index values).

## Data availability

Data will be made available on request to the corresponding author.

## Notes

### Competing Interest Statement

The authors have declared no competing interest.

### Summary of Updates

Sections 3.6. and 3.7 on the ene-reductase YqjM production in C. necator updated, activity tests included, Supplemental files updated.

