## Supplementary Information for "High-yield protein production in the chemolithoautotrophic bacterium *Cupriavidus necator* H16"

#### Supplementary Tables

**Table S1:** Strains used and created in this study.

| Strain | Genotype | Reference |
|---|---|---|
| *Cupriavidus necator* ΔRM | *C. necator* H16 ΔE6A55_RS00030 ΔE6A55_RS00035 ΔE6A55_RS00040 ΔE6A55_RS00045 | (Vajente et al., 2024) |
| *Cupriavidus necator* ΔnagR | *C. necator* ΔRM Δ*nagR* (E6A55_RS01580) | This study |
| *Cupriavidus necator* T7R | *C. necator* ΔRM Δ*nagR* (E6A55_RS01580) *rhaRS,* $P_{rha}$-RBS 2-*T7RNAP* (not codon optimized) | This study |
| *Cupriavidus necator* T7S | *C. necator* ΔRM Δ*nagR* (E6A55_RS01580) $P_{H16\_RS08125}$-*T7RNAP* (not codon optimized) | This study |
| *Cupriavidus necator* T7R2 | *C. necator* ΔRM *rhaRS,* $P_{rha}$-RBS 1-*T7RNAP* (codon harmonized) | This study |
| *Cupriavidus necator* T7RT | *C. necator* ΔRM *rhaRS,* $P_{rha}$-RBS 1-*T7RNAP* (codon harmonized) with SIBR insertion between G442 and L443 | This study |
| *Escherichia coli* DH5α | *F- Φ80lacZΔM15 Δ(lacZYA-argF) U169 recA1 endA1 hsdR17(rk-, mk+) phoA supE44 thi-1 gyrA96 relA1 λ-* | ThermoFisher Scientific |
| *Escherichia coli* S17-1 | (DSM 9079) *RP4−2(Km::Tn7,Tc::Mu-1) pro-82 recA1 endA1 thiE1 hsdR17 creC510* | DSMZ |
| *Escherichia coli* BL21(DE3) | *fhuA2 [lon] ompT gal (λ DE3) [dcm] ΔhsdS λ DE3 = λ sBamHIo ΔEcoRI-B int::(lacI::PlacUV5::T7 gene1) i21 Δnin5* | New England Biolabs |

**Table S2: Plasmids used in this study**

| Plasmid | Genotype | Reference |
|---|---|---|
| pMVRha | pBBR1 *rep*, *ori*; *Km*$^r$; RP4 *parABCDE*; *rhaRS*; $p_{J5[C2]}$-*spisPink*; RP4 *oriT* | (Vajente et al., 2025) |
| pMVRha-Ec | pMVRha; $p_{rha}$-*GFPmut3* (Ec) | This study |
| pMVRha-Cn | pMVRha; $p_{rha}$-*GFPmut3* (Cn) | This study |
| pMVRha-YqjM | pMVRha; $p_{rha}$-*yqjM* (Ec) | This study |
| pMVRha-YqjM-Cn | pMVRha; $p_{rha}$-*yqjM* (Cn) | This study |
| pET28a-YqjM | pET28a (pUC *ori*; *Km*$^r$; *lacI*) *yqjM* (Ec) | (Zhang et al., 2019) |
| pSEVA23g19[g1] | pBBR1 *rep*, ori; *Km*$^r$*; lacZ* | (Blázquez et al., 2022) |
| p1-Cn | pSEVA23g19[g1]; J23106-RBS 1-*GFPmut3* (Cn)-B1006 | This study |
| p2-Cn | pSEVA23g19[g1]; J23106-RBS 2-*GFPmut3* (Cn)-B1006 | This study |
| p3-Cn | pSEVA23g19[g1]; J23106-RBS 2-*GFPmut3* (Cn)-B1006 | This study |
| pT7α1-Ec | pSEVA23g19[g1]; T7 (consensus)-RBS 1-*GFPmut3* (Ec)-tT7hyb | This study |
| pT7α2-Ec | pSEVA23g19[g1]; T7 (consensus)-RBS 2-*GFPmut3* (Ec)-tT7hyb | This study |
| pT7α1-Cn | pSEVA23g19[g1]; T7 (consensus)-RBS 1-*GFPmut3* (Cn)-tT7hyb | This study |
| pT7α2-Cn | pSEVA23g19[g1]; T7 (consensus)-RBS 2-*GFPmut3* (Cn)-B1006 | This study |
| pT7α3-Cn | pSEVA23g19[g1]; T7 (consensus)-RBS 3-*GFPmut3* (Cn)-tT7hyb | This study |
| pT7β2-Cn | pSEVA23g19[g1]; T7 (variant 2)-RBS 1-*GFPmut3* (Cn)-tT7hyb | This study |
| pT7γ2-Cn | pSEVA23g19[g1]; T7 (variant 3)-RBS 1-*GFPmut3* (Cn)-tT7hyb | This study |

| | | |
|---|---|---|
| pT7β2-YqjM | pSEVA23g19[g1]; T7 (variant 2)-RBS 1-*yqjM* (Cn)-tT7hyb | This study |
| pLO3Mt | *$Km^r$; sacB*; RP4 *oriT*; pUC *ori*; silent mutation in restriction enzyme recognition site | (Vajente et al., 2025) |
| pLONagR | pLO3Mt; 1 kb homology regions near *nagR* for *nagR* deletion | This study |
| pLOT7R | pLO3Mt; 1 kb homology regions near *nagR* for *nagR* deletion; *rhaRS,* $P_{rha}$-RBS_A-*T7RNAP* (not codon optimized) | This study |
| pLOT7S | pLO3Mt; 1 kb homology regions near *nagR* for *nagR* deletion; $P_{H16\_RS08125}$-*T7RNAP* (not codon optimized) | This study |
| pLOT7R2 | pLO3Mt; 1 kb homology regions near *nagR*, without *nagR* deletion; *rhaRS,* $P_{rha}$-B0034m-*T7RNAP* (codon harmonized) | This study |
| pLOT7RT | pLO3Mt; 1 kb homology regions near *nagR*, without *nagR* deletion; *rhaRS,* $P_{rha}$-B0034m-*T7RNAP* (codon harmonized) with SIBR insertion between G442 and L443 | This study |

**Table S3: List of all genetic parts and fragments synthesized and used in this study.** Restriction enzyme recognition sites used for cloning are bold, restriction enzyme cutting sites are underlined.

| Genetic part | Source | Reference | Sequence |
|---|---|---|---|
| Promoter T7 α (consensus) | Synthesized as annealed oligonucleotides | (Jones et al., 2015) - Consensus promoter | **GGTCTC**TGGAGTAATACGACTCACTATAGGGGAATACTT**GAGACC** |
| Promoter T7 β (variant 2) | Synthesized as annealed oligonucleotides | (Jones et al., 2015) - C4 promoter | **GGTCTC**TGGAGTAATACGACTCACTATCAAGGAATACTT**GAGACC** |

| | | | |
|---|---|---|---|
| Promoter T7 γ (variant 3) | Synthesized as annealed oligonucleotides | (Conrad et al., 2020) | **GGTCTC**TGGAGTAATACGACTCACTATAGGGATAATTACTT**GAGACC** |
| Promoter J23106 | GoldenStandard library | (Blázquez et al., 2022) | **GGTCTC**AGGAGTTTACGGCTAGCTCAGTCCTAGGTATAGTGCTAGCTACTA**GAGACC** |
| RBS 1 (B0034m) | GoldenStandard library | (Blázquez et al., 2022) | **GGTCTC**TTACTAGAGAAAGAGGAGAAATACTAAATGT**GAGACC** |
| RBS 2 | Synthesized as annealed oligonucleotides | (Sydow et al., 2017) | **GGTCTC**TTACTAATGAACAATTCTTAAGAAGGAGATATACAAATGT**GAGACC** |
| RBS 3 | Synthesized as annealed oligonucleotides | This study | **GGTCTC**TTACTATAATTTTGTTTAACCAGAAGAAGGAGATATACAATGT**GAGACC** |
| CDS *GFPmut3* (Ec) | GoldenStandard library | (Blázquez et al., 2022) | **GGTCTC**TAATGATGCGTAAAGGAGAAGAACTTTTCACTGGAGTTGTCCCAATTCTTGTTGAATTAGATGGTGATGTTAATGGGCACAAATTTTCTGTCAGTGGAGAGGGTGAAGGTGATGCAACATACGGAAAACTTACCCTTAAATTTATTTGCACTACTGGAAAACTACCTGTTCCATGGCCAACACTTGTCACTACTTTCGGTTATGGTGTTCAATGCTTTGCGAGATACCCAGATCATATGAAACAGCATGACTTTTTCAAGAGTGCCATGCCCGAAGGTTATGTACAGGAAAGAACTATATTTTTCAAAGATGACGGGAACTACAAGACACGTGCTGAAGTCAAGTTTGAAGGTGATACCCTTGTTAATAGAATCGAGTTAAAAGGTATTGATTTTAAAGAAGATGGAAACATTCTTGGACACAAATTGGAATACAACTATAACTCACACAATGTATACATCATGGCAGACAAACAAAAGAATGGAATCAAAGTTAACTTCAAAATTAGACACAACATTGAAGATGGAAGCGTTCAACTAGCAGACCATTATCAACAAAATACTCCAATTGGCGATGGCCCTGTCCTTTTACCAGACAACCATTACCTGTCCACACAATCTGCCCTTTCGAAAGATCCCAACGAAAAGAGAGATCACATGGTCCTTCTTGAGTTTGTAACAGCTGCTGGGATTACACATGGCATGGATGAACTATACAAATAAGCTTT**GAGACC** |

| | | | |
|---|---|---|---|
| CDS *GFPmut3* (Cn) | Synthesized by Twist bioscience | This study | **GGTCTC**AAATGTCGAAGGGCGAAGAGCTGTTCACCGGGGTGGTACCCATCCTGGTGGAACTGGACGGCGATGTGAACGGCCATAAGTTCTCGGTGTCGGGCGAGGGCGAGGGCGACGCCACGTACGGCAAGCTGACGCTCAAGTTCATCTGCACCACGGGCAAGCTGCCCGTGCCATGGCCTACCTTGGTGACGACCTTCGGCTATGGAGTGCAGTGTTTCGCCCGCTATCCGGACCACATGAAGCAACATGACTTTTTCAAGTCGGCCATGCCGGAGGGCTACGTCCAGGAGCGCACCATCTTTTTCAAGGATGACGGGAATTACAAGACGCGAGCTGAAGTGAAGTTTGAAGGCGACACCCTGGTGAACCGGATAGAGCTGAAGGGCATCGACTTCAAGGAGGATGGCAACATCCTGGGTCACAAGCTGGAGTACAACTATAATTCCCACAACGTCTACATCATGGCCGACAAGCAGAAGAACGGCATCAAGGTCAACTTCAAGATCCGTCACAATATCGAGGACGGGTCCGTGCAGCTCGCGGACCACTATCAACAGAACACGCCGATCGGCGACGGCCCGGTGCTCCTGCCCGATAATCACTACCTTTCCACCCAGTCCGCATTGAGCAAGGACCCTAACGAAAAGCGTGACCATATGGTCCTGCTCGAGTTCGTCACTGCGGCCGGCATTACTCACGGCATGGACGAGCTGTACAAATGAGCTTA**GAGACC** |
| CDS *yqjM* (Ec) | Kindly provided by Prof. Frank Hollmann (PCR-amplified from pET28a-YqjM) | (Zhang et al., 2019) | **GGTCTC**TAATGGGCAGCAGCCATCATCATCATCATCACAGCAGCGGCCTGGTGCCGCGCGGCAGCCATATGGCCAGAAAATTATTTACACCTATTACAATTAAAGATATGACGTTAAAAAACCGCATTGTCATGTCGCCAATGTGCATGTATTCTTCTCATGAAAAGGACGGAAAATTAACACCGTTCCACATGGCACATTACATATCGCGCGCAATCGGCCAGGTCGGACTGATTATTGTAGAGGCGTCAGCGGTTAACCCTCAAGGACGAATCACTGACCAAGACTTAGGCATTTGGAGCGACGAGCATATTGAAGGCTTTGCAAAACTGACTGAGCAGGTCAAAGAACAAGGTTCAAAAATCGGCATTCAGCTTGCCCATGCCGGACGTAAAGCTGAGCTTGAAGGAGATATCTTCGCTCCATCGGCGATTGCGTTTGACGAACAATCAGCAACACCTGTAGAAATGTCAGCAGAAAAAGTAAAAGAAACGGTCCAGGAGTTCAAGCAAGCGGCTGCCCGCGCAAAAGAAGCCGGCTTTGATGTGATTGAAATTCATGCGGCGCACGGATATTTAATTCATGAATTTTTGTCTCCGCTTTCCAACCATCGAACAGATGAATATGGCGGCTCACCTGAAAACCGCTATCGTTTCTTGAGAGAGATCATTGATGAAGTCAAACAAGTATGGGACGGTCCTTTATTTGTCCGTGTATCTGCTTCTGACTACACTGATAAAGGCTTAG |

| | | | |
|---|---|---|---|
| | | | ACATTGCCGATCACATCGGTTTTGCAAAATGGATGAAGG<br>AGCAGGGTGTTGACTTAATTGACTGCAGCTCAGGCGCCCT<br>TGTTCACGCAGACATTAACGTATTCCCTGGCTATCAGGTC<br>AGCTTCGCTGAGAAAATCCGTGAACAGGCGGACATGGCT<br>ACTGGTGCCGTCGGCATGATTACAGACGGTTCAATGGCT<br>GAAGAAATTCTGCAAAACGGACGTGCCGACCTCATCTTT<br>ATCGGCAGAGAGCTTTTGCGGGATCCATTTTTTGCAAGA<br>ACTGCTGCGAAACAGCTCAATACAGAGATTCCGGCCCCTG<br>TTCAATACGAAAGAGGCTGGTAAGCTTT**GAGACC** |
| CDS *yqjM* (Cn) | Synthesized by Twist bioscience | This study | **GGTCTC**TAATGCACCATCACCACCATCACATGGCGCGCA<br>AGCTGTTCACCCCCATCACCATCAAGGACATGACGCTGAA<br>GAACCGCATCGTGATGTCCCCCATGTGCATGTACAGCAGC<br>CACGAGAAAGATGGCAAGCTGACCCCGTTTCATATGGCCC<br>ACTATATTTCCCGCGCCATCGGCCAGGTGGGCCTGATCAT<br>CGTCGAAGCCTCGGCCGTGAACCCCCAGGGCCGGATCACG<br>GATCAGGATCTGGGCATCTGGAGCGATGAACACATCGAG<br>GGCTTCGCCAAGCTGACGGAACAGGTGAAGGAGCAGGGG<br>TCGAAGATCGGCATCCAGCTGGCGCACGCGGGCCGCAAGG<br>CGGAACTGGAGGGCGACATCTTTGCGCCCTCCGCCATCGC<br>CTTCGATGAGCAGTCGGCCACCCCCGTCGAGATGTCGGCC<br>GAGAAGGTCAAGGAAACGGTGCAGGAATTTAAACAGGCC<br>GCGGCGCGCGCCAAGGAGGCGGGCTTCGACGTGATCGAG<br>ATCCACGCCGCCCATGGCTACCTGATCCACGAGTTCCTGA<br>GCCCGCTGTCCAACCACCGGACCGACGAGTACGGCGGCTC<br>GCCCGAGAACCGCTACCGCTTTCTGCGCGAAATCATCGAC<br>GAGGTGAAGCAGGTCTGGGATGGGCCCCTGTTCGTGCGC<br>GTCAGCGCGAGCGATTATACGGACAAGGGCCTGGATATC<br>GCGGACCATATCGGGTTCGCCAAGTGGATGAAAGAACAG<br>GGGGTGGATCTGATCGATTGCAGCTCGGGCGCGCTGGTGC<br>ATGCCGATATCAACGTCTTTCCCGGCTACCAGGTGAGCTT<br>TGCGGAAAAGATCCGCGAGCAGGCCGATATGGCGACGGG<br>GGCGGTGGGCATGATCACCGATGGGTCGATGGCGGAGGA<br>GATCCTGCAGAACGGCCGCGCGGATCTCATCTTCATCGGC<br>CGCGAACTGCTGCGGGACCCCTTCTTCGCCCGCACGGCGG<br>CCAAGCAGCTCAACACCGAAATCCCGGCGCCCGTGCAGTA<br>TGAGCGCGGCTGGTGAGCTTT**GAGACC** |

| | | | |
|---|---|---|---|
| Terminator tT7hyb | Synthesized as annealed oligonucleotides | (Calvopina-Chavez et al., 2022) - Terminator T7hyb1 | **GGTCTC**TGCTTAAACAGATAGGCCCTCTTCGGAGGGCCTATCTGTTTTTTTTTCGCTT**GAGACC** |
| Terminator B1006_GI | GoldenStandard library | (Blázquez et al., 2022) | **GGTCTC**TGCTTAAAAAAAAACCCCGCCCCTGACAGGGCGGGGTTTTTTTTCGCTT**GAGACC** |
| Terminator BBa_B0011 | Synthesized as annealed oligonucleotides | iGEM website | **GGTCTC**AAGAGAATATAAAAAGCCAGATTATTAATCCGGCTTTTTTATTATTTT**GAGACC** |
| Terminator BBa_B0014 | Synthesized by Twist bioscience | iGEM website | **GGTCTC**GGTAATCACACTGGCTCACCTTCGGGTGGGCCTTTCTGCGTTTATATACTAGAGAGAGAATATAAAAAGCCAGATTATTAATCCGGCTTTTTTATTATTTGCGAT**GAGACC** |
| Terminator BBa_B0015 + $P_{H16\_RS08125}$ | Synthesized by Twist bioscience | (Hanko et al., 2020) | **GGTCTC**GACCAGGCATCAAATAAAACGAAAGGCTCAGTCGAAAGACTGGGCCTTTCGTTTTATCTGTTGTTTGTCGGTGAACGCTCTCTACTAGAGTCACACTGGCTCACCTTCGGGTGGGCCTTTCTGCGTTTATAGGTGCCGTCTATTATTGATTTTATGAATGTTCATTATAGCCAGAATTCTATTGGTTAATTTTCCTCCGTGCACCATGATCCTTCCCATACGAAGAGACATAGCCGGAGACAT**GAGACC** |
| SIBR | Synthesized by Twist bioscience | (Della Valle et al., 2025) | **GGTCTC**AAGGTTAATTGAGGCCTGAGTATAAGGTGACTTATACTTGTAATCTATCTAAACGGGGAACCTCTCTAGTAGACAATCCCGTGCTAAATTGATACCAGCATCGTCTTGATGCCCTTGGCAGCATAAATGCCTAACGACTATCCCTTTGGGGAGTAGGGTCAAGTGACTCGAAACGATAGACAACTTGCTTTAACAAGTTGGAGATATAGTCTGCTCTGCATGGTGACATGCAGCTGGATATAATTCCGGGGTAAGATTAACGACCTTATCTGAACATAATGCTGCTCACGA**GAGACC** |
| T7 RNA polymerase (Ec) | PCR-amplified from *E. coli* | (Studier and | **GGTCTC**ATACAATGAACACGATTAACATCGCTAAGAACGACTTCTCTGACATCGAACTGGCTGCTATCCCGTTCAACACTCTGGCTGACCATTACGGTGAGCGTTTAGCTCGCGAACAG |

| | | | |
|---|---|---|---|
| | BL21(DE3) genomic DNA | Moffatt, 1986) | TTGGCCCTTGAGCATGAGTCTTACGAGATGGGTGAAGCA<br>CGCTTCCGCAAGATGTTTGAGCGTCAACTTAAAGCTGGT<br>GAGGTTGCGGATAACGCTGCCGCCAAGCCTCTCATCACTA<br>CCCTACTCCCTAAGATGATTGCACGCATCAACGACTGGTT<br>TGAGGAAGTGAAAGCTAAGCGCGGCAAGCGCCCGACAGC<br>CTTCCAGTTCCTGCAAGAAATCAAGCCGGAAGCCGTAGCG<br>TACATCACCATTAAGACCACTCTGGCTTGCCTAACCAGTG<br>CTGACAATACAACCGTTCAGGCTGTAGCAAGCGCAATCG<br>GTCGGGCCATTGAGGACGAGGCTCGCTTCGGTCGTATCCG<br>TGACCTTGAAGCTAAGCACTTCAAGAAAAACGTTGAGGA<br>ACAACTCAACAAGCGCGTAGGGCACGTCTACAAGAAAGC<br>ATTTATGCAAGTTGTCGAGGCTGACATGCTCTCTAAGGG<br>TCTACTCGGTGGCGAGGCGTGGTCTTCGTGGCATAAGGA<br>AGACTCTATTCATGTAGGAGTACGCTGCATCGAGATGCT<br>CATTGAGTCAACCGGAATGGTTAGCTTACACCGCCAAAA<br>TGCTGGCGTAGTAGGTCAAGACTCTGAGACTATCGAACT<br>CGCACCTGAATACGCTGAGGCTATCGCAACCCGTGCAGGT<br>GCGCTGGCTGGCATCTCTCCGATGTTCCAACCTTGCGTAG<br>TTCCTCCTAAGCCGTGGACTGGCATTACTGGTGGTGGCTA<br>TTGGGCTAACGGTCGTCGTCCTCTGGCGCTGGTGCGTACT<br>CACAGTAAGAAAGCACTGATGCGCTACGAAGACGTTTAC<br>ATGCCTGAGGTGTACAAAGCGATTAACATTGCGCAAAAC<br>ACCGCATGGAAAATCAACAAGAAAGTCCTAGCGGTCGCC<br>AACGTAATCACCAAGTGGAAGCATTGTCCGGTCGAGGAC<br>ATCCCTGCGATTGAGCGTGAAGAACTCCCGATGAAACCG<br>GAAGACATCGACATGAATCCTGAGGCTCTCACCGCGTGG<br>AAACGTGCTGCCGCTGCTGTGTACCGCAAGGACAAGGCTC<br>GCAAGTCTCGCCGTATCAGCCTTGAGTTCATGCTTGAGCA<br>AGCCAATAAGTTTGCTAACCATAAGGCCATCTGGTTCCCT<br>TACAACATGGACTGGCGCGGTCGTGTTTACGCTGTGTCA<br>ATGTTCAACCCGCAAGGTAACGATATGACCAAAGGACTG<br>CTTACGCTGGCGAAAGGTAAACCAATCGGTAAGGAAGGT<br>TACTACTGGCTGAAAATCCACGGTGCAAACTGTGCGGGT<br>GTCGATAAGGTTCCGTTCCCTGAGCGCATCAAGTTCATTG<br>AGGAAAACCACGAGAACATCATGGCTTGCGCTAAGTCTC<br>CACTGGAGAACACTTGGTGGGCTGAGCAAGATTCTCCGT<br>TCTGCTTCCTTGCGTTCTGCTTTGAGTACGCTGGGGTACA<br>GCACCACGGCCTGAGCTATAACTGCTCCCTTCCGCTGGCG |

| | | | |
|---|---|---|---|
| | | | TTTGACGGGTCTTGCTCTGGCATCCAGCACTTCTCCGCGA<br>TGCTCCGAGATGAGGTAGGTGGTCGCGCGGTTAACTTGC<br>TTCCTAGTGAAACCGTTCAGGACATCTACGGGATTGTTGC<br>TAAGAAAGTCAACGAGATTCTACAAGCAGACGCAATCAA<br>TGGGACCGATAACGAAGTAGTTACCGTGACCGATGAGAA<br>CACTGGTGAAATCTCTGAGAAAGTCAAGCTGGGCACTAA<br>GGCACTGGCTGGTCAATGGCTGGCTTACGGTGTTACTCGC<br>AGTGTGACTAAGCGTTCAGTCATGACGCTGGCTTACGGG<br>TCCAAAGAGTTCGGCTTCCGTCAACAAGTGCTGGAAGAT<br>ACCATTCAGCCAGCTATTGATTCCGGCAAGGGTCTGATGT<br>TCACTCAGCCGAATCAGGCTGCTGGATACATGGCTAAGCT<br>GATTTGGGAATCTGTGAGCGTGACGGTGGTAGCTGCGGT<br>TGAAGCAATGAACTGGCTTAAGTCTGCTGCTAAGCTGCT<br>GGCTGCTGAGGTCAAAGATAAGAAGACTGGAGAGATTCT<br>TCGCAAGCGTTGCGCTGTGCATTGGGTAACTCCTGATGGT<br>TTCCCTGTGTGGCAGGAATACAAGAAGCCTATTCAGACG<br>CGCTTGAACCTGATGTTCCTCGGTCAGTTCCGCTTACAGC<br>CTACCATTAACACCAACAAAGATAGCGAGATTGATGCAC<br>ACAAACAGGAGTCTGGTATCGCTCCTAACTTTGTACACA<br>GCCAAGACGGTAGCCACCTTCGTAAGACTGTAGTGTGGG<br>CACACGAGAAGTACGGAATCGAATCTTTTGCACTGATTC<br>ACGACTCCTTCGGTACCATTCCGGCTGACGCTGCGAACCT<br>GTTCAAAGCAGTGCGCGAAACTATGGTTGACACATATGA<br>GTCTTGTGATGTACTGGCTGATTTCTACGACCAGTTCGCT<br>GACCAGTTGCACGAGTCTCAATTGGACAAAATGCCAGCA<br>CTTCCGGCTAAAGGTAACTTGAACCTCCGTGACATCTTAG<br>AGTCGGACTTCGCGTTCGCGTAAT**GAGACC** |
| T7 RNA polymerase (Cn) + terminator BBa_B0014 | Synthesized by Twist bioscience | This study, iGEM website | **GGTCTC**AAATGAACACGATCAACATCGCCAAGAACGACT<br>TCTCGGACATCGAACTGGCCGCCATCCCCTTCAACACCCT<br>GGCCGACCATTACGGCGAGCGCCTCGCCCGCGAACAGCTC<br>GCACTGGAGCATGAGTCGTACGAGATGGGCGAAGCACGC<br>TTCCGCAAGATGTTTGAGCGCCAGCTGAAAGCCGGCGAG<br>GTGGCAGATAACGCCGCAGCAAAGCCGCTCATCACCACCC<br>TCCTCCCGAAGATGATCGCACGCATCAACGACTGGTTTGA<br>GGAAGTGAAAGCCAAGCGCGGGAAGCGCCCCACGGCATT<br>CCAGTTCCTGCAGGAAATCAAGCCCGAAGCAGTGGCATAC<br>ATCACCATCAAGACCACCCTGGCCTGCCTCACCTCCGCCG<br>ACAATACGACCGTGCAGGCCGTGGCATCCGCAATCGGCCG |

| | | | |
|---|---|---|---|
| | | | TGCAATCGAGGACGAGGCCCGCTTCGGCCGCATCCGCGAC<br>CTGGAAGCCAAGCACTTCAAGAAAAACGTGGAGGAACAG<br>CTCAACAAGCGCGTGGGGCACGTCTACAAGAAAGCATTT<br>ATGCAGGTGGTCGAGGCCGACATGCTCTCGAAGGGCCTCC<br>TCGGCGGGGAGGCATGGTCGTCATGGCATAAGGAAGACT<br>CGATCCATGTGGGGGTGCGCTGCATCGAGATGCTCATCGA<br>GTCCACCGGGATGGTGTCCCTCCACCGCCAGAATGCCGGG<br>GTGGTGGGCCAGGACTCGGAAACCATCGAACTCGCACCGG<br>AATACGCCGAGGCCATCGCAACCCGCGCAGGCGCACTGGC<br>CGGGATCTCGCCCATGTTCCAGCCGTGCGTGGTGCCGCCG<br>AAGCCCTGGACCGGGATCACCGGCGGCGGGTATTGGGCCA<br>ACGGCCGCCGCCCGCTGGCACTGGTGCGCACCCACTCCAA<br>GAAAGCACTGATGCGCTACGAAGACGTGTACATGCCGGA<br>GGTGTACAAAGCAATCAACATCGCACAGAACACGGCATG<br>GAAAATCAACAAGAAAGTCCTCGCAGTCGCAAACGTGAT<br>CACCAAGTGGAAGCATTGCCCCGTCGAGGACATCCCGGCA<br>ATCGAGCGCGAAGAACTCCCCATGAAACCCGAAGACATC<br>GACATGAATCCGGAGGCCCTCACCGCATGGAAACGCGCCG<br>CAGCCGCCGTGTACCGCAAGGACAAGGCCCGCAAGTCGCG<br>CCGCATCTCCCTGGAGTTCATGCTGGAGCAGGCAAATAAG<br>TTTGCCAACCATAAGGCAATCTGGTTCCCGTACAACATGG<br>ACTGGCGCGGCCGCGTGTACGCCGTGTCCATGTTCAACCC<br>CCAGGGCAACGATATGACCAAAGGGCTGCTGACGCTGGC<br>AAAAGGCAAACCCATCGGCAAGGAAGGCTACTACTGGCT<br>GAAAATCCACGGCGCAAACTGCGCAGGCGTCGATAAGGT<br>GCCCTTCCCGGAGCGCATCAAGTTCATCGAGGAAAACCAC<br>GAGAACATCATGGCCTGCGCCAAGTCGCCCCTGGAGAACA<br>CCTGGTGGGCCGAGCAGGATTCGCCCTTCTGCTTCCTGGC<br>ATTCTGCTTTGAGTACGCCGGGGTGCAGCACCACGGGCTG<br>TCCTATAACTGCTCCCTGCCCCTGGCATTTGACGGGTCGT<br>GCTCGGGGATCCAGCACTTCTCCGCAATGCTCCGGGATGA<br>GGTGGGCGGCCGCGCAGTGAACCTCCTGCCGTCCGAAACC<br>GTGCAGGACATCTACGGGATCGTGGCCAAGAAAGTCAAC<br>GAGATCCTCCAGGCAGACGCAATCAATGGGACCGATAAC<br>GAAGTGGTGACCGTGACCGATGAGAACACCGGCGAAATC<br>TCGGAGAAAGTCAAGCTGGGGACCAAGGCACTGGCCGGC<br>CAGTGGCTGGCCTACGGCGTGACCCGCTCCGTGACCAAGC<br>GCTCCGTCATGACGCTGGCCTACGGGTCCAAAGAGTTCGG |

| | | | |
|---|---|---|---|
| | | | GTTCCGCCAGCAGGTGCTGGAAGATACCATCCAGCCCGCC<br>ATCGATTCCGGGAAGGGCCTGATGTTCACCCAGCCCAATC<br>AGGCCGCCGGGTACATGGCCAAGCTGATCTGGGAATCGG<br>TGTCCGTGACGGTGGTGGCCGCAGTGGAAGCAATGAACT<br>GGCTGAAGTCGGCCGCCAAGCTGCTGGCCGCCGAGGTCAA<br>AGATAAGAAGACCGGGGAGATCCTGCGCAAGCGCTGCGC<br>CGTGCATTGGGTGACCCCGGATGGCTTCCCGGTGTGGCAG<br>GAATACAAGAAGCCGATCCAGACGCGCCTCAACCTGATG<br>TTCCTCGGCCAGTTCCGCCTCCAGCCGACCATCAACACCA<br>ACAAAGATTCCGAGATCGATGCACACAAACAGGAGTCGG<br>GCATCGCCCCGAACTTTGTGCACTCCCAGGACGGCTCCCA<br>CCTGCGCAAGACCGTGGTGTGGGCACACGAGAAGTACGG<br>GATCGAATCGTTTGCACTGATCCACGACTCCTTCGGCACC<br>ATCCCCGCCGACGCCGCAAACCTGTTCAAAGCAGTGCGCG<br>AAACCATGGTGGACACGTATGAGTCGTGCGATGTGCTGG<br>CCGATTTCTACGACCAGTTCGCCGACCAGCTCCACGAGTC<br>GCAGCTCGACAAAATGCCCGCACTGCCCGCCAAAGGCAAC<br>CTCAACCTCCGCGACATCCTCGAGTCAGACTTCGCATTCG<br>CGTAATCACACTGGCTCACCTTCGGGTGGGCCTTTCTGCG<br>TTTATATACTAGAGAGAGAATATAAAAAGCCAGATTATT<br>AATCCGGCTTTTTTATTATTTGCGAATCCTCGTAGGTACC<br>ACTAT**GAGACC** |
| T7 RNA polymerase (Cn) + SIBR + terminator BBa_B0014 | Synthesized by Twist bioscience | This study, iGEM website, (Della Valle et al., 2025) | **GGTCTC**AAATGAACACGATCAACATCGCCAAGAACGACT<br>TCTCGGACATCGAACTGGCCGCCATCCCCTTCAACACCCT<br>GGCCGACCATTACGGCGAGCGCCTCGCCCGCGAACAGCTC<br>GCACTGGAGCATGAGTCGTACGAGATGGGCGAAGCACGC<br>TTCCGCAAGATGTTTGAGCGCCAGCTGAAAGCCGGCGAG<br>GTGGCAGATAACGCCGCAGCAAAGCCGCTCATCACCACCC<br>TCCTCCCGAAGATGATCGCACGCATCAACGACTGGTTTGA<br>GGAAGTGAAAGCCAAGCGCGGGAAGCGCCCCACGGCATT<br>CCAGTTCCTGCAGGAAATCAAGCCCGAAGCAGTGGCATAC<br>ATCACCATCAAGACCACCCTGGCCTGCCTCACCTCCGCCG<br>ACAATACGACCGTGCAGGCCGTGGCATCCGCAATCGGCCG<br>TGCAATCGAGGACGAGGCCCGCTTCGGCCGCATCCGCGAC<br>CTGGAAGCCAAGCACTTCAAGAAAAACGTGGAGGAACAG<br>CTCAACAAGCGCGTGGGGCACGTCTACAAGAAAGCATTT<br>ATGCAGGTGGTCGAGGCCGACATGCTCTCGAAGGGCCTCC<br>TCGGCGGGGAGGCATGGTCGTCATGGCATAAGGAAGACT |

| | | | |
|---|---|---|---|
| | | | CGATCCATGTGGGGGTGCGCTGCATCGAGATGCTCATCGA<br>GTCCACCGGGATGGTGTCCCTCCACCGCCAGAATGCCGGG<br>GTGGTGGGCCAGGACTCGGAAACCATCGAACTCGCACCGG<br>AATACGCCGAGGCCATCGCAACCCGCGCAGGCGCACTGGC<br>CGGGATCTCGCCCATGTTCCAGCCGTGCGTGGTGCCGCCG<br>AAGCCCTGGACCGGGATCACCGGCGGCGGGTATTGGGCCA<br>ACGGCCGCCGCCCGCTGGCACTGGTGCGCACCCACTCCAA<br>GAAAGCACTGATGCGCTACGAAGACGTGTACATGCCGGA<br>GGTGTACAAAGCAATCAACATCGCACAGAACACGGCATG<br>GAAAATCAACAAGAAAGTCCTCGCAGTCGCAAACGTGAT<br>CACCAAGTGGAAGCATTGCCCCGTCGAGGACATCCCGGCA<br>ATCGAGCGCGAAGAACTCCCCATGAAACCCGAAGACATC<br>GACATGAATCCGGAGGCCCTCACCGCATGGAAACGCGCCG<br>CAGCCGCCGTGTACCGCAAGGACAAGGCCCGCAAGTCGCG<br>CCGCATCTCCCTGGAGTTCATGCTGGAGCAGGCAAATAAG<br>TTTGCCAACCATAAGGCAATCTGGTTCCCGTACAACATGG<br>ACTGGCGCGGCCGCGTGTACGCCGTGTCCATGTTCAACCC<br>CCAGGGCAACGATATGACCAAAGGTTAATTGAGGCCTGA<br>GTATAAGGTGACTTATACTTGTAATCTATCTAAACGGGG<br>AACCTCTCTAGTAGACAATCCCGTGCTAAATTGATACCAG<br>CATCGTCTTGATGCCCTTGGCAGCATAAATGCCTAACGAC<br>TATCCCTTTGGGGAGTAGGGTCAAGTGACTCGAAACGAT<br>AGACAACTTGCTTTAACAAGTTGGAGATATAGTCTGCTC<br>TGCATGGTGACATGCAGCTGGATATAATTCCGGGGTAAG<br>ATTAACGACCTTATCTGAACATAATGCTGCTCACGCTGGC<br>AAAAGGCAAACCCATCGGCAAGGAAGGCTACTACTGGCT<br>GAAAATCCACGGCGCAAACTGCGCAGGCGTCGATAAGGT<br>GCCCTTCCCGGAGCGCATCAAGTTCATCGAGGAAAACCAC<br>GAGAACATCATGGCCTGCGCCAAGTCGCCCCTGGAGAACA<br>CCTGGTGGGCCGAGCAGGATTCGCCCTTCTGCTTCCTGGC<br>ATTCTGCTTTGAGTACGCCGGGGTGCAGCACCACGGGCTG<br>TCCTATAACTGCTCCCTGCCCCTGGCATTTGACGGGTCGT<br>GCTCGGGGATCCAGCACTTCTCCGCAATGCTCCGGGATGA<br>GGTGGGCGGCCGCGCAGTGAACCTCCTGCCGTCCGAAACC<br>GTGCAGGACATCTACGGGATCGTGGCCAAGAAAGTCAAC<br>GAGATCCTCCAGGCAGACGCAATCAATGGGACCGATAAC<br>GAAGTGGTGACCGTGACCGATGAGAACACCGGCGAAATC<br>TCGGAGAAAGTCAAGCTGGGGACCAAGGCACTGGCCGGC |

| | | | |
|---|---|---|---|
| | | | CAGTGGCTGGCCTACGGCGTGACCCGCTCCGTGACCAAGC<br>GCTCCGTCATGACGCTGGCCTACGGGTCCAAAGAGTTCGG<br>GTTCCGCCAGCAGGTGCTGGAAGATACCATCCAGCCCGCC<br>ATCGATTCCGGGAAGGGCCTGATGTTCACCCAGCCCAATC<br>AGGCCGCCGGGTACATGGCCAAGCTGATCTGGGAATCGG<br>TGTCCGTGACGGTGGTGGCCGCAGTGGAAGCAATGAACT<br>GGCTGAAGTCGGCCGCCAAGCTGCTGGCCGCCGAGGTCAA<br>AGATAAGAAGACCGGGGAGATCCTGCGCAAGCGCTGCGC<br>CGTGCATTGGGTGACCCCGGATGGCTTCCCGGTGTGGCAG<br>GAATACAAGAAGCCGATCCAGACGCGCCTCAACCTGATG<br>TTCCTCGGCCAGTTCCGCCTCCAGCCGACCATCAACACCA<br>ACAAAGATTCCGAGATCGATGCACACAAACAGGAGTCGG<br>GCATCGCCCCGAACTTTGTGCACTCCCAGGACGGCTCCCA<br>CCTGCGCAAGACCGTGGTGTGGGCACACGAGAAGTACGG<br>GATCGAATCGTTTGCACTGATCCACGACTCCTTCGGCACC<br>ATCCCCGCCGACGCCGCAAACCTGTTCAAAGCAGTGCGCG<br>AAACCATGGTGGACACGTATGAGTCGTGCGATGTGCTGG<br>CCGATTTCTACGACCAGTTCGCCGACCAGCTCCACGAGTC<br>GCAGCTCGACAAAATGCCCGCACTGCCCGCCAAAGGCAAC<br>CTCAACCTCCGCGACATCCTCGAGTCAGACTTCGCATTCG<br>CGTAATCACACTGGCTCACCTTCGGGTGGGCCTTTCTGCG<br>TTTATATACTAGAGAGAGAATATAAAAAGCCAGATTATT<br>AATCCGGCTTTTTTATTATTTGCGAATCCTCGTAGGTACC<br>ACTAT**GAGACC** |

**Table S4: Codon usage tables of *E. coli* BL21 (NCBI: GCF_013166975) and *C. necator* H16 (NCBI: GCF_004798725) calculated using cusp (EMBOSS).**

| **Codon** | **AA** | **Frequency *E. coli* [%]** | **Frequency *C. necator* [%]** |
|---|---|---|---|
| GCA | A | 21.3 | 8.9 |
| GCC | | 26.8 | 47.4 |
| GCG | | 35.6 | 39.1 |
| GCT | | 16.2 | 4.6 |
| TGC | C | 55.4 | 90.5 |
| TGT | | 44.6 | 9.5 |
| GAC | D | 37.3 | 73.8 |
| GAT | | 62.7 | 26.2 |
| GAA | E | 69.3 | 40.2 |

| | | | |
|---|---|---|---|
| GAG | | 30.7 | 59.8 |
| TTC | F | 42.7 | 81.5 |
| TTT | | 57.3 | 18.5 |
| GGA | G | 10.8 | 4.6 |
| GGC | | 40.4 | 75.3 |
| GGG | | 15 | 12.6 |
| GGT | | 33.8 | 7.5 |
| CAC | H | 43.3 | 63.9 |
| CAT | | 56.7 | 36.1 |
| ATA | I | 7.4 | 1.9 |
| ATC | | 41.9 | 86.1 |
| ATT | | 50.8 | 12 |
| AAA | K | 76.6 | 10.3 |
| AAG | | 23.4 | 89.7 |
| CTA | L | 3.6 | 1.2 |
| CTC | | 10.5 | 13.8 |
| CTG | | 49.7 | 74 |
| CTT | | 10.3 | 4.8 |
| TTA | | 13 | 0.3 |
| TTG | | 12.8 | 5.9 |
| ATG | M | 100 | 100 |
| AAC | N | 55 | 76.2 |
| AAT | | 45 | 23.8 |
| CCA | P | 19.1 | 5.9 |
| CCC | | 12.3 | 27.4 |
| CCG | | 52.7 | 60.8 |
| CCT | | 15.8 | 5.9 |
| CAA | Q | 34.8 | 14 |
| CAG | | 65.2 | 86 |
| AGA | R | 3.7 | 0.9 |
| AGG | | 2.2 | 3.8 |
| CGA | | 6.5 | 2.8 |
| CGC | | 40 | 67.3 |
| CGG | | 9.7 | 17.5 |
| CGT | | 37.9 | 7.6 |
| AGC | S | 27.6 | 35 |
| AGT | | 15 | 3.5 |

| | | | |
|---|---|---|---|
| TCA | | 12.4 | 3.9 |
| TCC | | 14.9 | 18 |
| TCG | | 15.6 | 36.9 |
| TCT | | 14.6 | 2.7 |
| ACA | T | 13 | 4.1 |
| ACC | | 43.3 | 57 |
| ACG | | 26.9 | 34.3 |
| ACT | | 16.8 | 4.6 |
| GTA | V | 15.6 | 4 |
| GTC | | 21.4 | 32.2 |
| GTG | | 37.1 | 58.8 |
| GTT | | 26 | 5 |
| TGG | W | 100 | 100 |
| TAC | Y | 42.9 | 69.8 |
| TAT | | 57.1 | 30.2 |
| TAA | * | 61.7 | 16.7 |
| TAG | | 7.7 | 16 |
| TGA | | 30.6 | 67.3 |

**Table S5: Table of calculated Codon Adaptation Index values (CAI) for all genes expressed in this study.**

| **Gene** | **CAI *E. coli* BL21 (NCBI: GCF_013166975)** | **CAI *C. necator* H16 (NCBI: GCF_004798725)** |
|---|---|---|
| *GFPmut3* (Ec) | 0.594 | 0.23 |
| *GFPmut3* (Cn) | 0.674 | 0.662 |
| *T7 RNA polymerase* (Ec) | 0.656 | 0.38 |
| *T7 RNA polymerase* (Cn) | 0.715 | 0.697 |
| *yqjM* (Ec) | 0.665 | 0.304 |
| *yqjM* (Cn) | 0.757 | 0.789 |

### Supplementary Figures

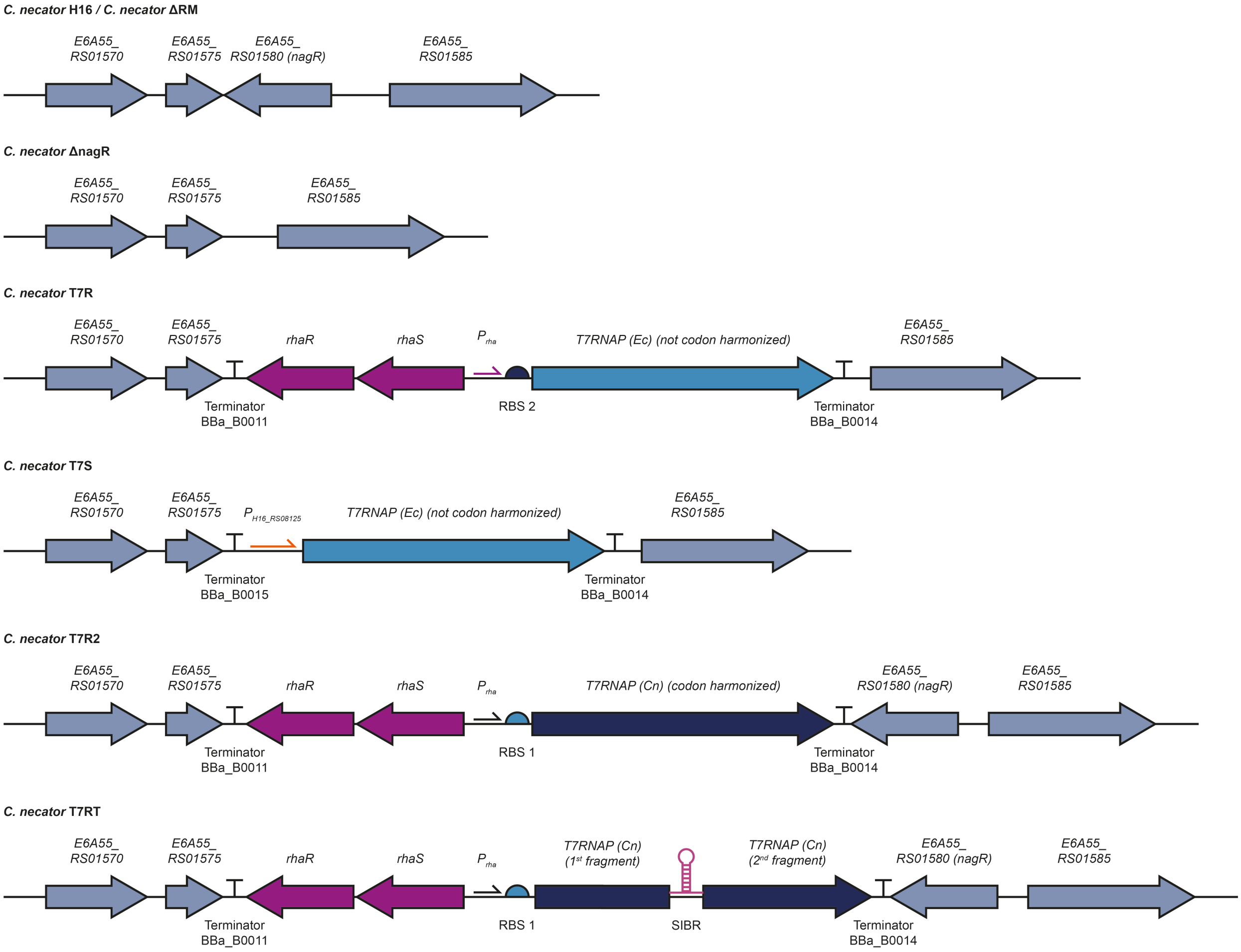


**Figure S1:** Schematic representation of the genotype of the strains created in this manuscript.

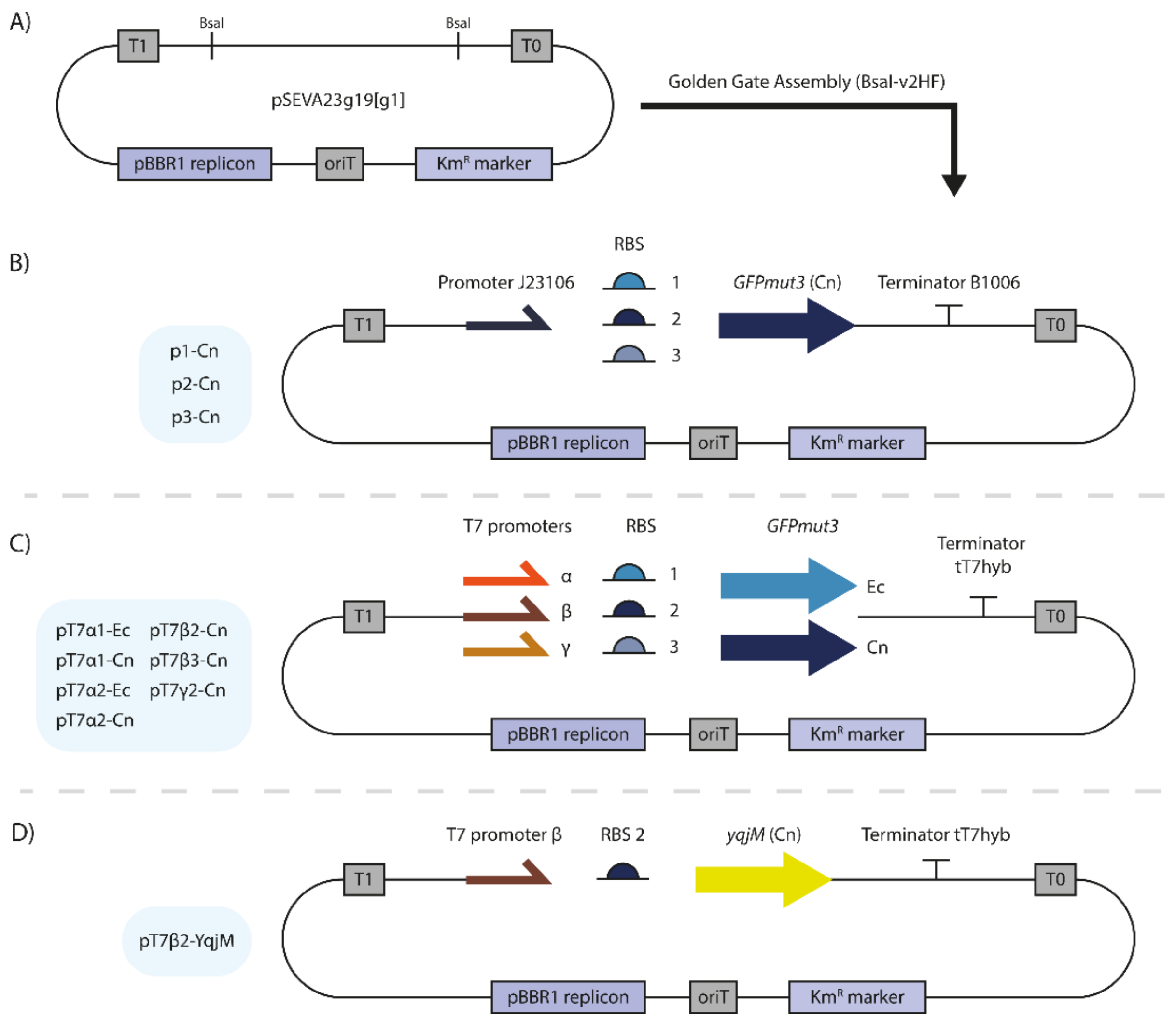


**Figure S2: Cloning diagram of plasmids derived from pSEVA23g19[g1];** A) Plasmid map of acceptor plasmid pSEVA23g19[g1]; B) Plasmids used for RBS strength analysis (Figure 3B, 5B). C) Plasmids used for genetic parts characterization (Figure 3, 4, 5); C) Plasmid used for *yqjM* expression (Figure 6).

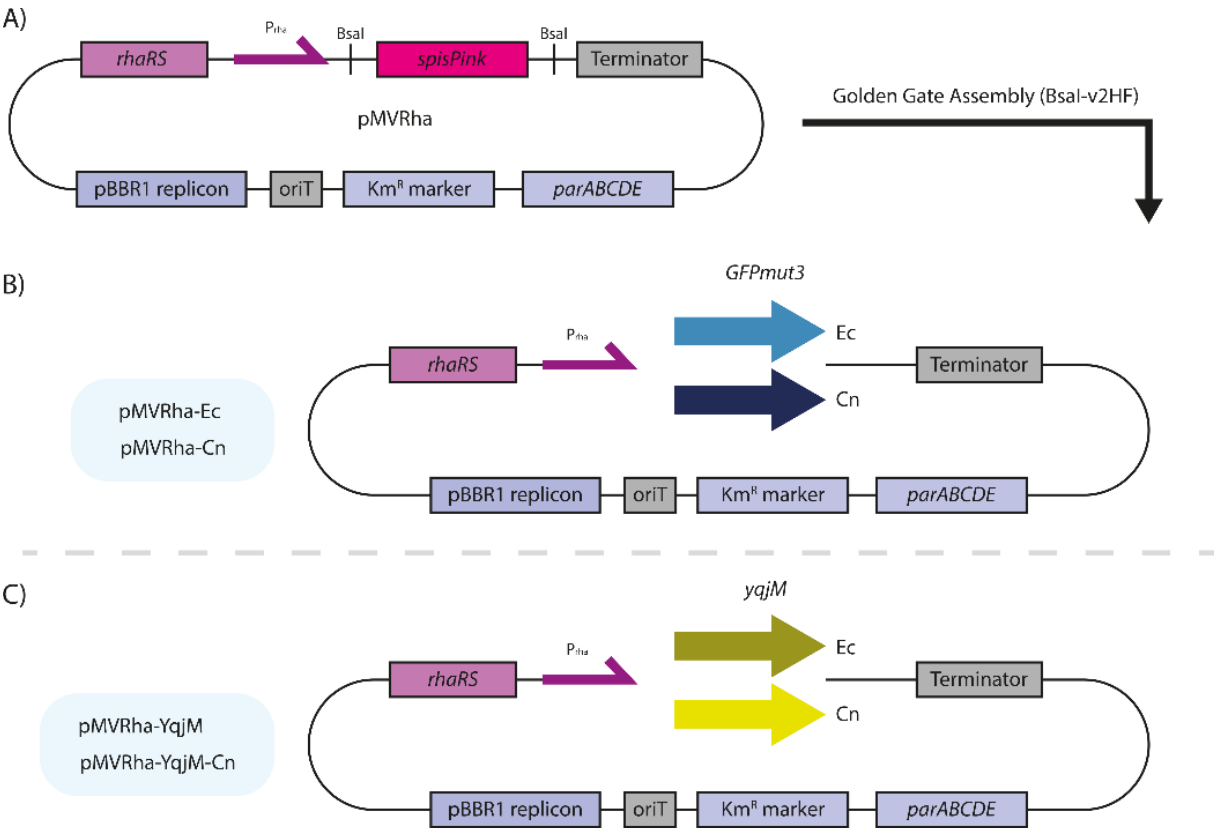


**Figure S3: Cloning diagram of plasmids derived from pMVRha;** A) Plasmid map of acceptor plasmid pMVRha; B) Plasmids used for *GFPmut3* expression (Figure 3, 4); C) Plasmids used for *yqjM* expression (Figure 6).

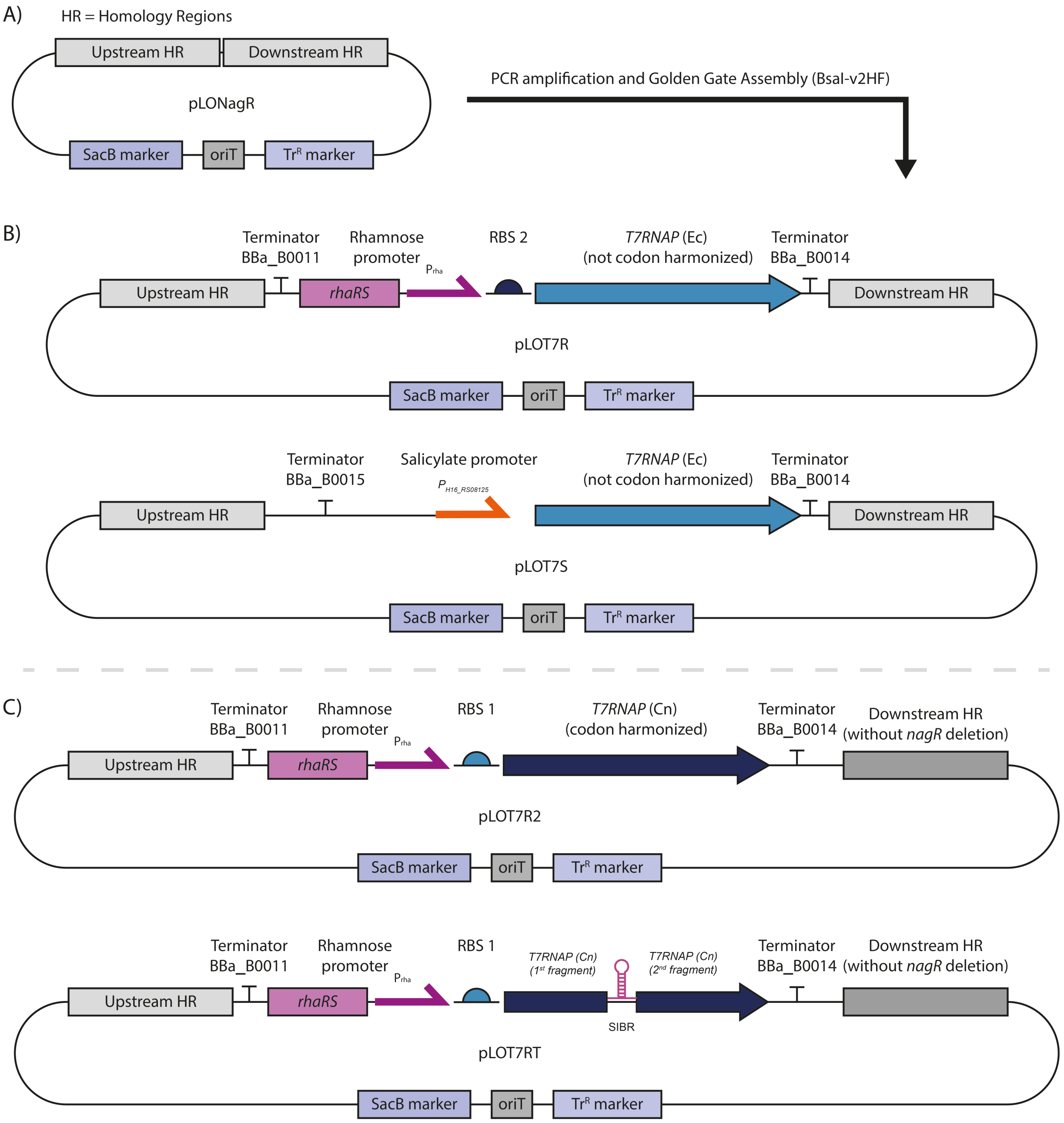


**Figure S4: Cloning diagram of plasmids derived from pLONagR, used for genomic integration in *C. necator*;** A) Plasmid map of acceptor plasmid pLONagR, derived from pLO3Mt; B) Plasmids used for the creation of strains *C. necator* T7R, *C. necator* T7S (Figure 1, S1); C) Plasmids used for the creation of strains *C. necator* T7R2, *C. necator* T7RT (Figure 4, S1).

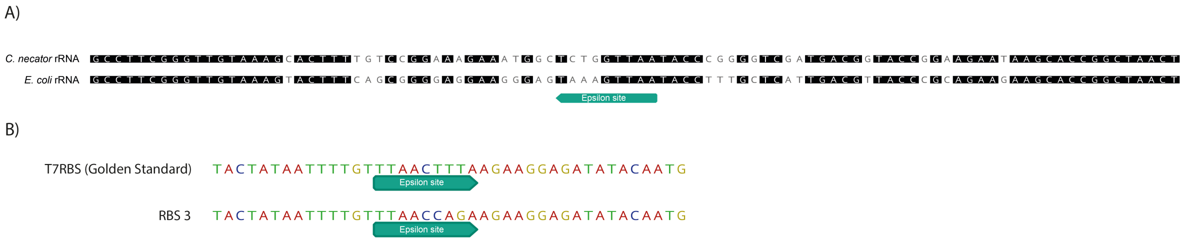


**Figure S5: Design of RBS 3 for *C. necator*.** A) Zoom-in of the alignment between rRNA sequences of *E. coli* and *C. necator*, with the original Epsilon sequence highlighted. B) Alignment of the pET RBS sequence (T7RBS from the Golden Standard library) and the newly designed RBS 3.

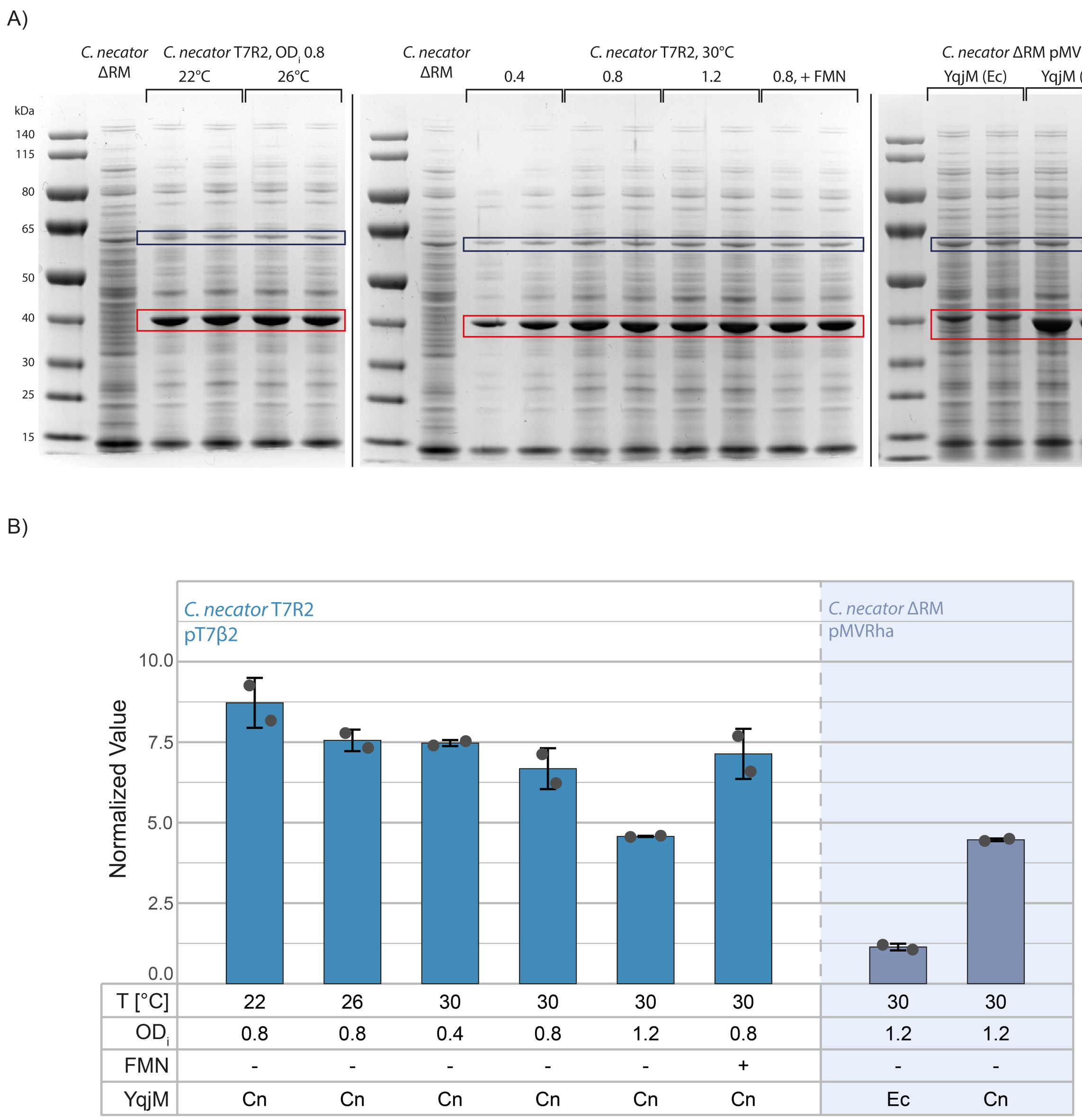


**Figure S6: Relative quantification of soluble YqjM produced by *C. necator* in different conditions, measured from SDS-PAGE.** The YqjM band intensity (red square) was divided by another protein band's intensity (blue square) to normalize the data. The assumption is that the intensity of this second band is constants in all conditions tested. Band intensities were quantified using the Gel Analysis tools from Fiji(Schindelin et al., 2012). T[°C]: incubation temperature after induction; OD$_i$: induction $OD_{600}$; FMN: in one experiment (+), FMN was supplemented to the culture at a final concentration of 1 µM; YqjM: two sequences with different codon usages were used (Ec, Cn) (see Table S5 for CAI and Table S3 for sequences). Arithmetic means and standard deviations of two biological replicates are reported.

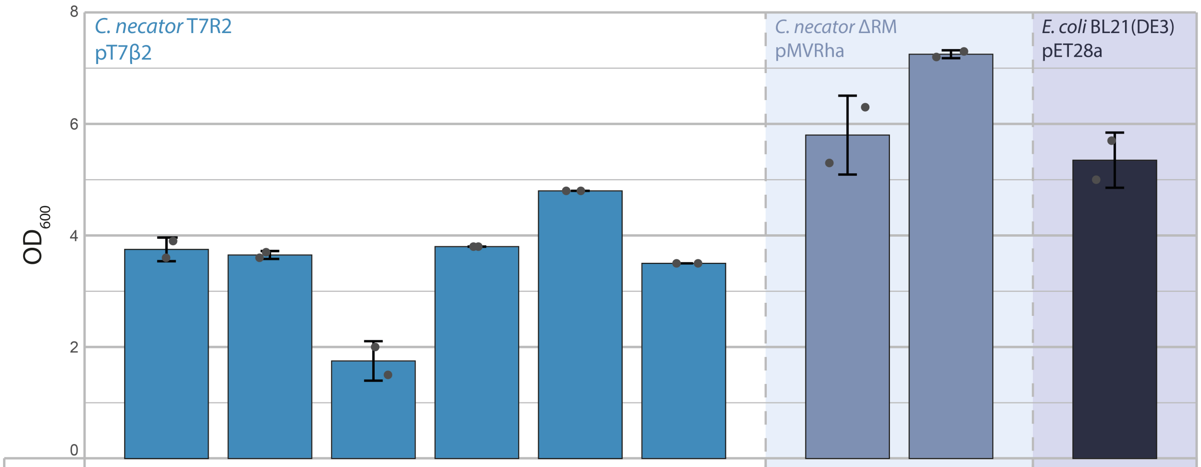


| T [°C] | 22 | 26 | 30 | 30 | 30 | 30 | 30 | 30 | 20 |
|---|---|---|---|---|---|---|---|---|---|
| $OD_i$ | 0.8 | 0.8 | 0.4 | 0.8 | 1.2 | 0.8 | 1.2 | 1.2 | 0.6 |
| FMN | - | - | - | - | - | + | - | - | - |
| YqjM | Cn | Cn | Cn | Cn | Cn | Cn | Ec | Cn | Ec |

**Figure S7: $OD_{600}$ reached by the strains analyzed in Figure 6.** $OD_{600}$ reached by the strains at the moment of harvest, measured using a cuvette spectrophotometer. T[°C]: incubation temperature after induction; $OD_i$: induction $OD_{600}$; FMN: in one experiment (+), FMN was supplemented to the culture at a final concentration of 1 µM; YqjM: two sequences with different codon usages were used (Ec, Cn) (see Table S5 for CAI and Table S3 for sequences). Arithmetic means and standard deviations of two biological replicates are reported.

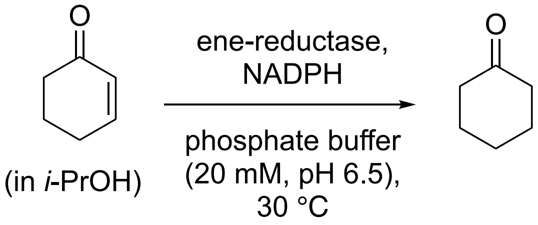


**Figure S8: Model reaction for YqjM activity assay.** Specific activities were recorded spectrophotometrically from the decrease of NADPH in YqjM catalyzed reduction of 2-cyclohexen-1-one. Reaction conditions: 30 °C; 20 mM potassium phosphate buffer (pH 6.5); 0.25 mM NADPH; 0.5 μM YqjM, 1 mM 2-cyclohexen-1-one in *i*-PrOH (100 mM final concentration).

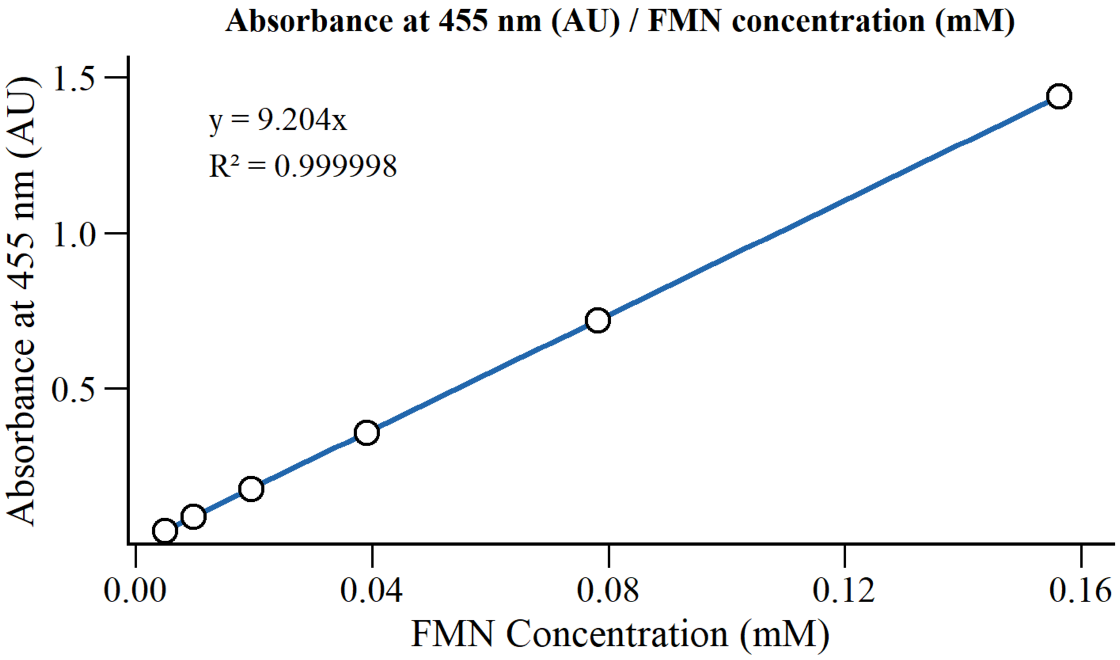


**Figure S9: FMN calibration curve.** Riboflavin 5′-monophosphate sodium salt was dissolved in storage buffer (20 mM potassium phosphate, pH 6.5) and serially diluted. The absorbance at 445 nm was measured using a spectrophotometer (Jasco V-650, quartz cuvette). Storage buffer was used to blank the instrument. Three different dilution series were measured.

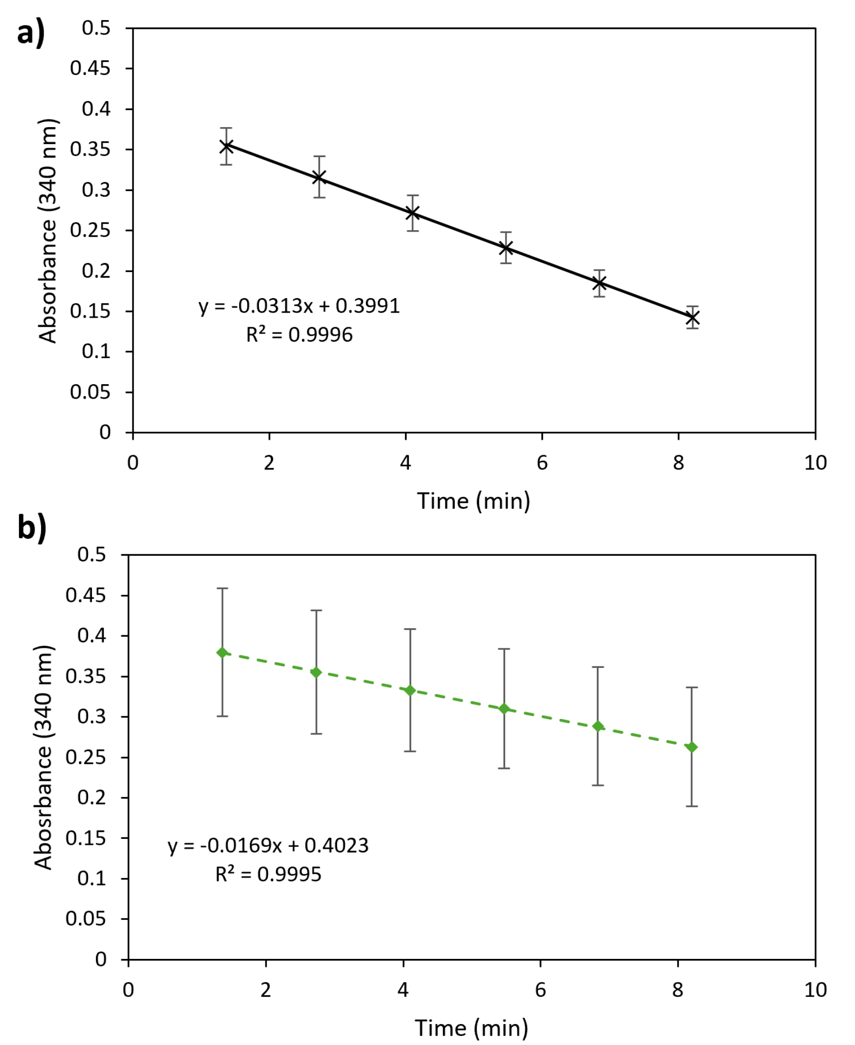


**Figure S9: Representative NADPH depletion curves obtained in the YqjM activity assay.** (a) Total NADPH depletion in the presence of substrate. (b) Background NADPH oxidation in the absence of substrate, with molecular oxygen serving as the sole electron acceptor. Specific activities were determined from the mean of triplicate measurements after subtraction of the background oxidation rate from the total NADPH depletion rate.
